# Sex differences in nociceptor translatomes contribute to divergent prostaglandin signaling in male and female mice

**DOI:** 10.1101/2020.07.31.231753

**Authors:** Diana Tavares-Ferreira, Pradipta R. Ray, Ishwarya Sankaranarayanan, Galo L. Mejia, Andi Wangzhou, Stephanie Shiers, Ruta Uttarkar, Salim Megat, Paulino Barragan-Iglesias, Gregory Dussor, Armen N. Akopian, Theodore J. Price

## Abstract

**Background:** There are clinically relevant sex differences in acute and chronic pain mechanisms, but we are only beginning to understand their mechanistic basis. Transcriptome analyses of rodent whole dorsal root ganglion (DRG) have revealed sex differences, mostly in immune cells. We examined the transcriptome and translatome of the mouse DRG with the goal of identifying sex differences.

**Methods:** We used Translating Ribosome Affinity Purification (TRAP) sequencing and behavioral pharmacology to test the hypothesis that nociceptor (Nav1.8 expressing neurons) translatomes would differ by sex.

**Results:** We found 66 genes whose mRNA were sex-differentially bound to nociceptor ribosomes. Many of these genes have known neuronal functions but have not been explored in sex differences in pain. We focused on *Ptgds*, which was increased in female mice. The mRNA encodes the prostaglandin D_2_ (PGD_2_) synthesizing enzyme. We observed increased Ptgds protein and PGD_2_ in female mouse DRG. The Ptgds inhibitor AT-56 caused intense pain behaviors in male mice but was only effective at high doses in females. Conversely, female mice responded more robustly to another major prostaglandin, PGE_2_, than did male mice. Ptgds protein expression was also higher in female cortical neurons, suggesting DRG findings may be generalizable to other nervous system structures.

**Conclusions:** Nociceptor TRAP sequencing (TRAP-seq) reveals unexpected sex differences in one of the oldest known nociceptive signaling molecule families, the prostaglandins. Our results demonstrate that translatome analysis reveals physiologically relevant sex differences important for fundamental protective behaviors driven by nociceptors.

## INTRODUCTION

For decades, studies in the neuroscience field have been done almost entirely on male animals (1). However, there are physiological and molecular differences in the peripheral and central nervous systems between males and females. Many neurological disorders have been shown to have different incidence proportion, age of onset, symptoms, and response to treatment between males and females. Schizophrenia tends to develop at an earlier age in men (2, 3); Parkinson’s disease is twice as common in men than women (4) and it has sex differences in symptoms and response to treatment (5). On the other hand, major depressive disorder (6), anxiety disorders (7), and Alzheimer’s disease (8) affect more women than men. There are also sex differences in pain syndromes. Neuropathic pain, osteoarthritis, migraine, and fibromyalgia are more frequently reported in women than in men (9–13). A common thread in all of these neurological disorders is that they are poorly treated by current therapies. The exclusion of one of the sexes in pre-clinical studies has likely been detrimental to the success of translational research. The National Institutes of Health has responded to this issue by mandating consideration of sex as a biological variable in funding proposals (14).

As noted above, there are clear sex differences in pain mechanisms, yet, we are only beginning to understand how these differences emerge (15). Nociceptors of the dorsal root (DRG) and trigeminal ganglion are the neurons that send nociceptive information to the brain and are a possible source of mechanistic diversity that causes sex differences in pain. A previous study suggested that there are some sex differences in sensory neuron transcriptomes (16). However, transcription and translation are not directly coupled in eukaryotes, so there can be important divergences between transcriptomes (the cellular RNA profile) and translatomes (the subset of the transcriptome that is bound to ribosomes for translation) in cells (17). An example in nociceptors is the similar transcription of the prolactin receptor (*Prlr*) in male and female nociceptors, but the female-selective localized translation of the *Prlr* mRNA (18). This sex difference in *Prlr* mRNA translation causes an important sex difference in prolactin-evoked pain responses (18–20).

The primary goal of our study was to identify differences in translatome between male and female nociceptors using Translating Ribosome Affinity Purification (TRAP) methodology (21–23). We demonstrate that there are specific mRNAs that are differentially bound by ribosomes (hence likely differentially translated) in neurons of the DRG between males and females. Notably, we identify differentially translated mRNAs encoding proteins that function in similar pathways between male and female nociceptors. This suggests that different proteins may drive similar functions in male and female neurons. One of the differentially translated mRNAs identified using our TRAP approach was *Ptgds* (Prostaglandin D-synthase). Ptgds is an abundant enzyme in neuronal cells that converts PGH_2_ to PGD_2_. We find that it is up-regulated in female neurons. Consistent with this, we observed significant differences in behavioral responses between males and females when inhibiting this enzyme. We also noted sex differences in the response to PGE_2_. Our use of TRAP technology to delve into sex differences in nociceptor translatomes reveals a fundamental difference in how male and female mice respond to one of the oldest known families of nociceptive signaling molecules, the prostaglandins. (24).

## METHODS AND MATERIALS

Detailed methods are provided in *Supplemental Methods*. All animal procedures were approved by the Institutional Animal Care and Use Committee at University of Texas at Dallas. DRGs from Nav1.8-TRAP male and female mice were quickly dissected and homogenized using Precellys® Minilys Tissue Homogenizer. An aliquot of the lysate was saved for use as INPUT (IN; bulk RNA-sequencing), and the remaining was used for immunoprecipitation (IP; TRAP-sequencing) by incubating the lysate with protein G-coated Dynabeads (Invitrogen) bound to anti-GFP antibodies for 3 h at 4°C. RNA was eluted from all samples using the Direct-zol kit (Zymo Research) and cDNA libraries were prepared with total RNA Gold library preparation (Illumina). After standardizing the amount of cDNA, the libraries were sequenced on Illumina NextSeq500 sequencing machine with 75-bp single-end reads. Reads were then mapped against the reference genome and transcriptome (Gencode vM16 and GRCm38.p5) using STAR v2.2.1 (25). Relative abundances in Transcripts Per Million (TPM) for each gene of each sample were quantified by Stringtie v1.3.5 (26). Downstream analyses were restricted to protein-coding genes and excluding mitochondrial chromosome genes. For each expressed coding gene, we report log_2_ fold change, Bhattacharyya coefficient (BC) (27) and strictly standardized mean difference (SSMD) (28, 29). We used immunohistochemistry, ELISA assay and behavioral tests to understand the consequences of sex differences in *Ptgds* expression.

## RESULTS

Nav1.8^cre^ mice were crossed with Rosa26^fs-TRAP^ (30) in order to create mice expressing eGFP fused to the ribosomal L10a protein in Nav1.8-positive neurons (Nav1.8-TRAP mice) (**Figure 1A**). The specificity of the TRAP approach has been previously characterized, where it was shown that eGFP-RPL10a is expressed in sensory neurons in the dorsal root ganglion (DRG) including most nociceptors (22, 23). We used Nav1.8-TRAP mice to characterize the translatome of male and female DRG neurons by immunoprecipitating actively translating ribosomes and purifying the associated mRNA (**Figure 1B**). We sequenced bulk RNA (INPUT or IN in later plots), correspondent to total RNA from whole DRGs, and immunoprecipitated (IP) mRNA, correspondent to mRNA bound to GFP-tagged ribosomes in Nav1.8-positive neurons. This approach allowed us to characterize differences at the steady-state transcriptional and active translational levels between female and male DRG.

**Figure 1:**
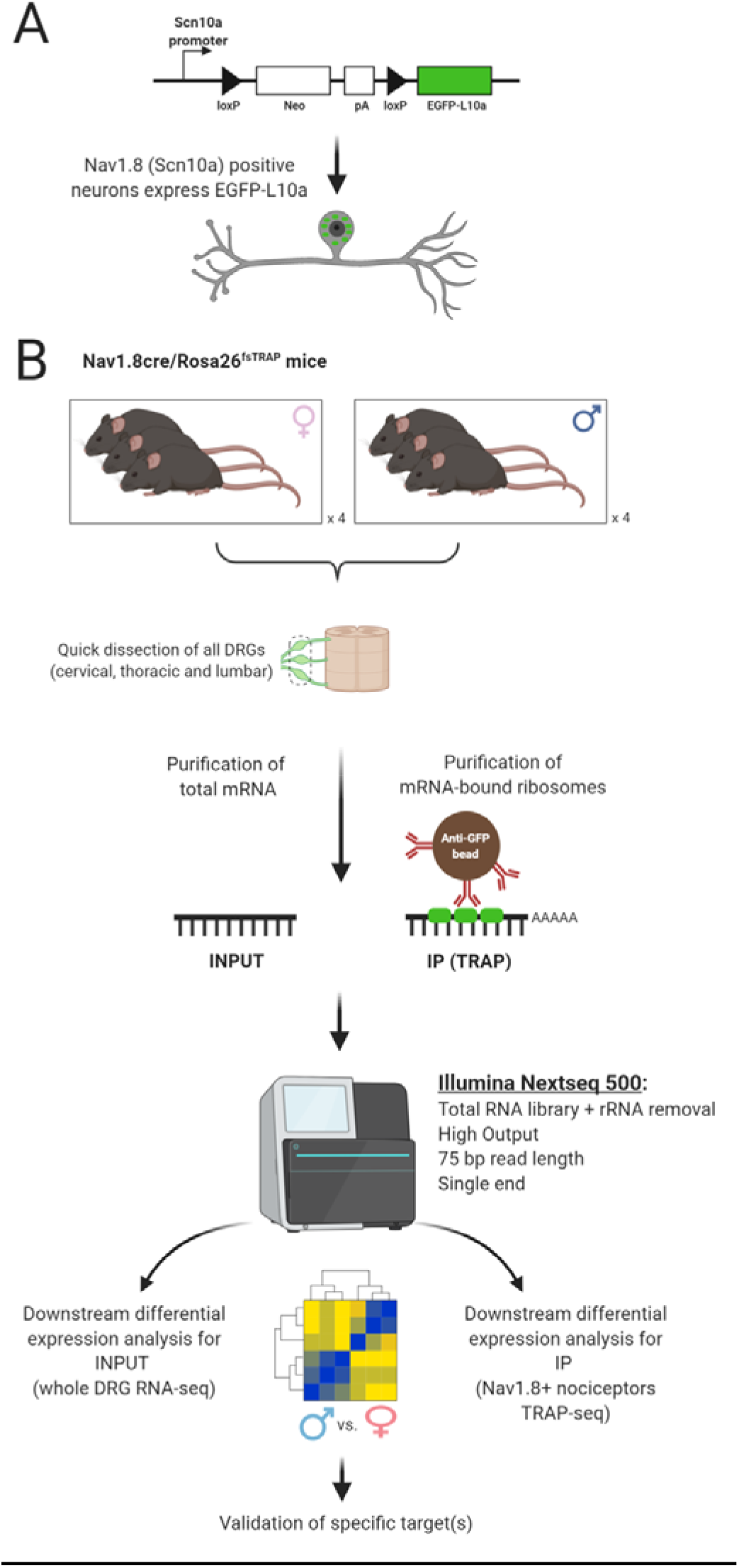
Outline of workflow for TRAP sequencing to reveal sex differences in nociceptor translatomes. **A**, eGFP-L10a protein is expressed in Nav1.8-positive nociceptors. **B**, Schematic representation of the methodology shows dissection of all DRGs (cervical, thoracic and lumbar) from Nav1.8cre/Rosa26^fsTRAP^ mice followed by isolation of total RNA (INPUT), and mRNA-bound to the ribosome (IP) using anti-eGFP-coated beads; Samples were sequenced and processed for downstream analysis of differentially expressed genes as shown.

Heatmap of the correlation coefficients for coding gene TPMs of each biological replicate showed a clear separation between TRAP-seq and bulk RNA-seq as expected given the different cell populations from which the cDNA library is constructed (**Figure 2A**). Consequently, in our hierarchical clustering analysis using these correlation coefficients, we find two distinct clusters of the bulk RNA-seq and TRAP-seq molecular profiles (**Figure 2B**). Subclusters within each cluster are very similar to each other and do not segregate by sex, showing that whole-transcriptome molecular profiles in each assay type (RNA-seq and TRAP-seq) are consistent across sexes for the DRG. All biological replicates for each sex in the input and IP samples showed high correlation coefficients across gene TPMs, suggesting high reproducibility across experiments (**Figure 2C**).

**Figure 2:**
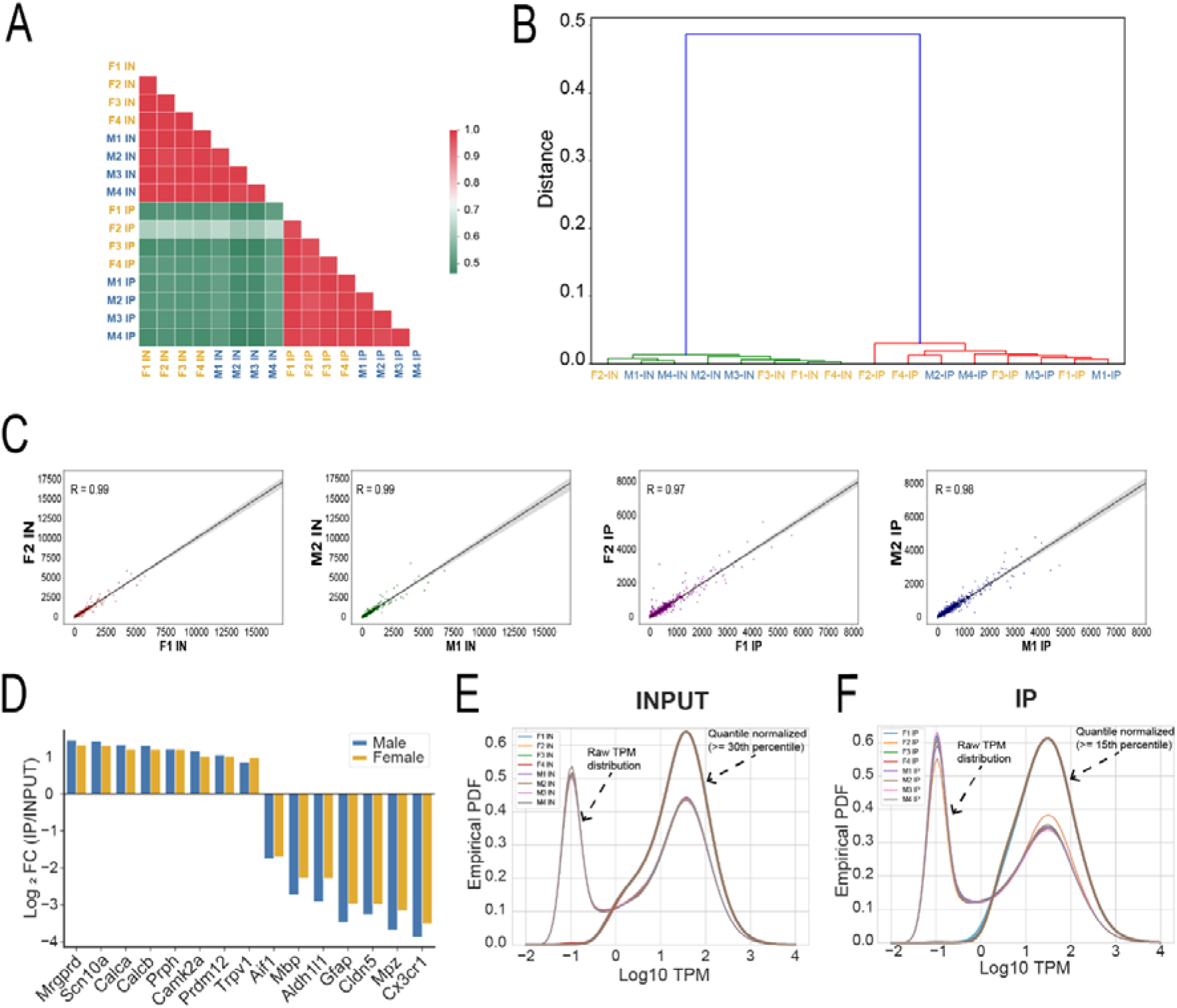
Nociceptor TRAP sequencing quality control. **A**, Hierarchical clustering analysis and **B**, Heatmap of the correlation coefficient show clear separation between TRAP-seq and bulk RNA-seq. However, we did not observe a clear distinction between male and female samples. **C**, Linear correlation plots shows high correlation coefficients of gene TPMs within biological replicates for the INPUT and IP fractions (shown for 2 replicates in each sex and assay), suggesting high reproducibility between replicates. **D**, Neuronal markers were enriched in IP fractions, such as *Calca* (encoding CGRP) or *Prph* (peripherin), while glial markers such as *Mbp*, *Mpz*, and *Gfap* were depleted (based on fold changes of median TPMs in each assay). **E, F**, The empirically estimated probability density of the raw TPMs and quantile normalized TPM (qTPM) distributions for the INPUT and IP fractions of all samples are shown. For INPUT samples, TPM distributions are shown for all coding genes and qTPMs shown for systematically transcriptome-expressed genes. For IP samples, TPM distributions were plotted for all coding genes, and qTPMs are shown for systematically translatome-expressed genes.

Because TRAP-seq purifies translating mRNAs in a cell type-specific manner, we tested the specificity of our approach using a group of control genes. We analyzed a subset of genes known to be enriched in specific cell populations in the DRG and verified that neuronal mRNAs, such as *Calca*, *Trpv1, Scn10a, Prph*, had enriched relative abundance in IP fraction. In contrast, non-neuronal genes such as glial markers (*Mpz*, *Mbp, Gfap*) were depleted in IP samples (**Figure 2D**).

Percentile ranks were calculated (**Suppl. File 1 Tabs 1A, 2A**) for gene expression levels (in TPM) for each RNA-seq and TRAP-seq sample. Based on these order statistics, we conservatively determined a set of 15,072 genes (>= 30th percentile) that were consistently detected in at least one sex in the RNA-seq samples, and out of those, 12,542 genes (>= 15th percentile) that were consistently detected in at least one sex in the TRAP-seq samples. These numbers are consistent with previous mouse DRG RNA-seq and TRAP-seq studies (31, 32).

For each biological replicate in INPUT and IP, we plotted the empirical probability densities of coding gene TPMs and noted a distinctly bimodal distribution for genes that are consistently detected versus those that are lowly expressed or undetected in each assay (**Figure 2E, F**). The TPM expression levels were finally quantile normalized (represented as qTPMs) in order to correct for sequencing depth and, thus, ensure comparability between samples.

To determine differentially expressed (DE) genes, we calculated the log2-fold change of TPM across sexes, and two related statistics - strictly standardized mean difference (SSMD) of TPM percentile ranks (28, 29) and Bhattacharyya coefficient (BC) of qTPMs (27) between sexes for quantifying the effect size and controlling for within-group variability. (**Suppl. File 1 Tabs 1A-B** for INPUT; **Suppl. File 1 Tabs 2A-B** for IP). For stringency, we required DE genes to have | log2-fold change | > 0.41 (corresponding to fold change > 1.33), | SSMD | > 0.9, and BD < 0.5. We plotted the SSMD values against the Fold Change (Log_2_ scale) for the autosomal genes for INPUT and IP (**Figures 3A, D**).

**Figure 3:**
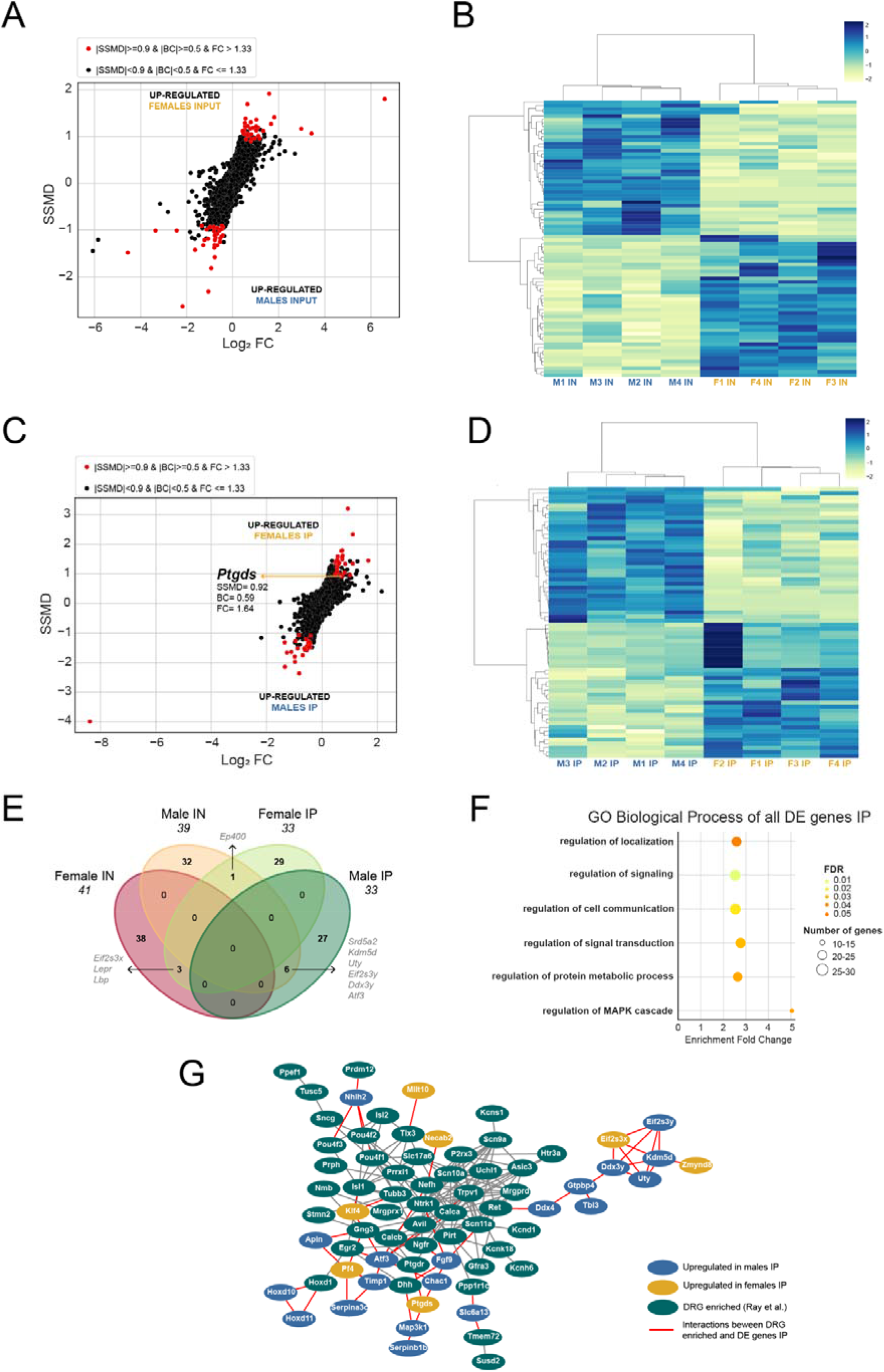
Differentially translated mRNAs in male and female DRG nociceptors. **A**, Dual-flashlight plot of INPUT samples showing SSMD and Log_2_ fold change values for all autosomal genes on or above the 30th percentile. **B**, Heatmap shows the z-scores of the differentially expressed genes in INPUT (IN) samples. Labels represent sex and biological replicate number. **C**, Dual-flashlight plot of IP samples showing SSMD and Log_2_ fold change values for all autosomal genes on or above the 15th percentile. **D**, Heatmap shows the z-scores of the differentially translated mRNAs in IP samples. **E**, Venn diagram comparing the genes identified as differentially expressed. There were few overlaps between INPUT and IP autosomal genes. **F**, GO terms enriched for all genes differentially expressed in IP. **G**, Network of interactions between genes differentially translated between males and females in IP and genes enriched in DRG neurons (Network generated using String database and Cytoscape).

We found a total of 80 genes for INPUT (transcriptome) (**Figure 3A, B**; **Suppl. File 2**) and 66 genes for IP (translatome) (**Figure 3C, D**; **Suppl. File 3**) that were DE between sexes. Given that we used mice that had not been experimentally manipulated, we anticipated finding a relatively small number of genes using both approaches. Interestingly, we did not observe a substantial overlap between genes DE in INPUT and IP, except for sex-chromosomal genes such as *Eif2s3x, Eif2s3y, Uty* and *Kdm5d* and a few autosomal genes (**Figure 3E**). We found that *Lepr* (Leptin receptor) and *Lbp* (Lipopolysaccharide binding protein) were up-regulated in both INPUT and IP, in females. *Ep400* (E1A Binding Protein P400) was up-regulated in male INPUT and female IP, highlighting discrepancies between transcriptome and translatome. *Atf3* (Activating transcription factor 3), a known nerve injury marker (33), was up-regulated in both INPUT and IP in males. Since *Atf3* can be induced by minor injuries, like scratches (34), this may be explained by more frequent fighting in male cages.

Next, we conducted gene-set enrichment analysis for DE genes using GO Enrichment Analysis resource PANTHER (35). We did not find any statistically significant GO terms for DE genes in INPUT. In contrast, we identified 6 GO terms (Biological Process) statistically significant (FDR < 0.05) for genes DE in IP (**Figure 3F**). The majority of the genes DE in IP were involved in the regulation of cell communication and signaling (**Figure 3G**). We identified several genes (e.g., *Sfrp4*, *Sema6a*) that encode for membrane proteins involved in cell signaling. Genes such as *Fbln1*, an extracellular matrix structural protein, and *Fgf9*, a growth factor, are involved in cell to cell communication. Genes such as *Map3k1* and *Mapk1ip1* are involved in the regulation of MAPK cascade. These findings suggest that there may be sex differences in proteins that regulate fundamental signaling pathways in male and female nociceptors. We expanded our functional analysis by manually curating relevant information regarding DE genes for INPUT and IP (**Suppl. Tables 1 and 2** contain detailed information).

Within the DE genes in the INPUT we identified several transcription factors such as *Hoxd4* in females and *Foxd3* (known to be glially expressed) in males. We also noted different immune-related genes identified in male and female DRG INPUT, consistent with previous work in mice (36, 37) and humans (38). These included *Cxcl16* (Chemokine (c-x-c motif) ligand 16), a T-cell signaling molecule, and *Jchain* (Immunoglobulin joining chain), which were up-regulated in female INPUT. *Cd276* (Cd276 antigen) was up-regulated in male INPUT and plays a role in inflammatory responses by regulating cytokine production and T cell receptor signaling.

In the IP fraction, as expected, the identified DE genes were neuronally enriched in expression when compared to the INPUT fraction (**Suppl. Tables 1 and 2**). In the female translatome (IP) we found several up-regulated DE genes that are involved in neuronal functions such as *Pcdha8* (Protocadherin alpha 8), *Zmynd8* (Zinc finger, mynd-type containing 8) and *Slc6a13* (Solute carrier family 6 (neurotransmitter transporter, GABA), member 13). In the male translatome, genes with known neuronal functions such as *Chka* (Choline kinase alpha) and *Sema6a* (Sema domain, transmembrane domain (tm), and cytoplasmic domain, (semaphorin) 6a) were found to be up-regulated. Several enriched gene-set categories were similar between males and females, but these were driven by different genes, suggesting that unique genes may control similar functions in male and female nociceptors.

Since translation efficiency can be controlled by sequence elements within the mRNA (39, 40), we examined whether there were motifs in the 5′ untranslated regions (UTRs) or 3’ UTRs of DE mRNAs in IP. We found three enriched motifs in the 3’ UTRs of the genes up-regulated in males (**Figure 4A**) and one enriched motif in the 3’ UTRs of the genes up-regulated in females (**Figure 4B**). The 3’ UTR of a gene is known to influence the localization, degradation, and translation efficiency of an mRNA (6,7). The identified motifs are involved in neuron differentiation and migration, cell communication, and signal transduction, in agreement with the biological functions of DE genes identified in our IP fractions. We did not find any enriched motifs in the 5’ UTRs of male or female DE mRNAs.

**Figure 4:**
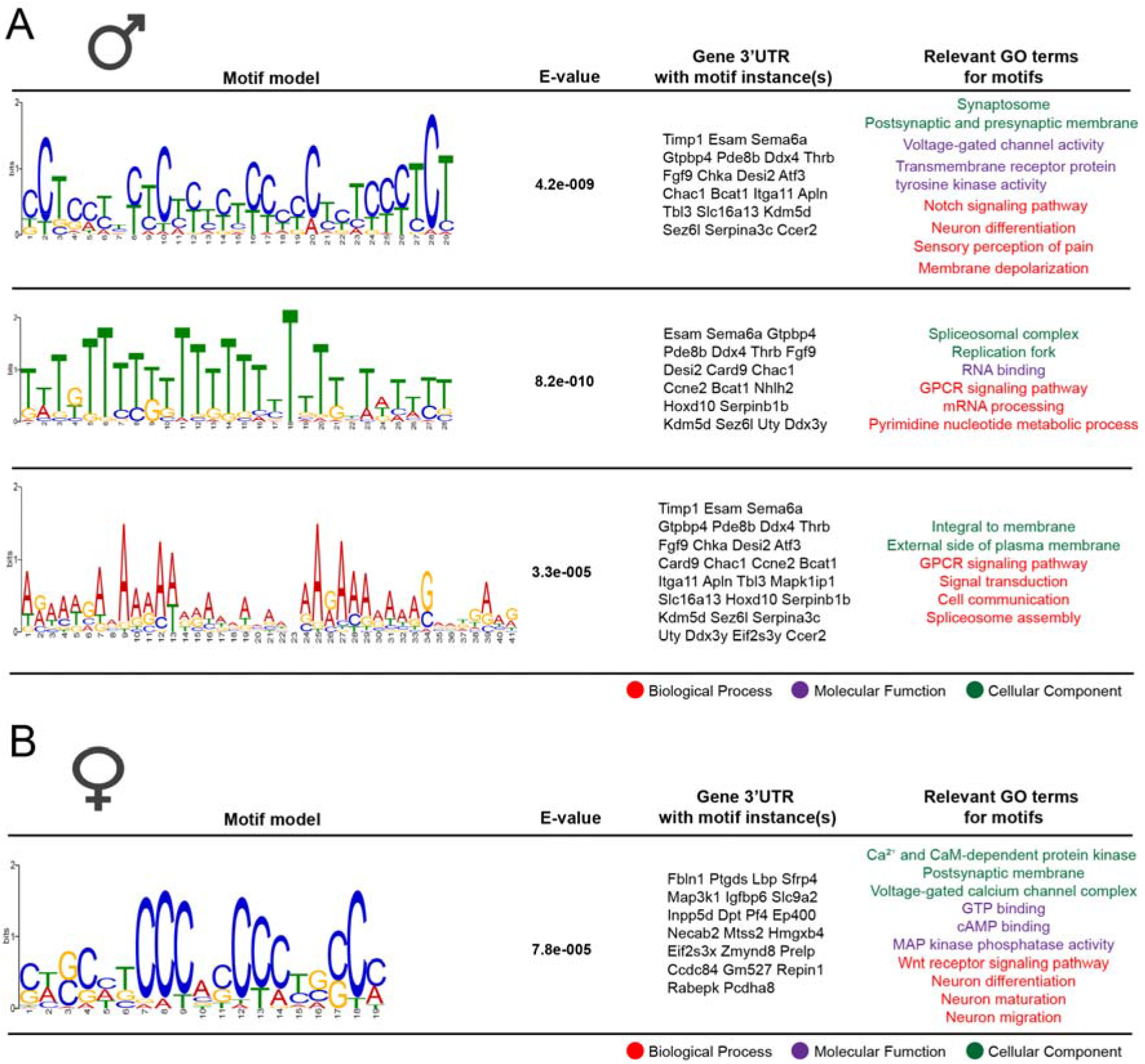
Enriched motifs identified in the 5’ or 3’ UTRs of mRNAs differentially translated in males or females. The motif analysis was conducted on the list of up-regulated mRNAs in both male and female IP fractions. **A**, We found 3 motifs significantly enriched in the 3’ UTR of up-regulated male mRNAs compared to the female mRNAs. **B**, We identified one motif significantly enriched in the 3’ UTR of up-regulated female mRNAs compared to the male mRNAs.

Next, we proceeded to validate our TRAP-seq approach by linking sex differences in the nociceptor active translatome to functional differences in expression and/or behavior. We decided to focus on a gene that was up-regulated in the female IP: Prostaglandin-D2 synthase (*Ptgds*). Ptgds catalyzes the conversion of prostaglandin H_2_ (PGH_2_) to prostaglandin D2 (PGD_2_) (**Figure 5A**), which is known to be the most abundant prostaglandin in the brain (41, 42) and regulates nociception, sleep and temperature homeostasis (43–53). Prostaglandins are among the most widely studied of pain inducing molecules and many drugs are currently marketed as analgesics that target this class of molecules (54–56). We reasoned that previously unknown sex differences in prostaglandin signaling could have a dramatic impact on further our understanding of sex differences in pain.

**Figure 5:**
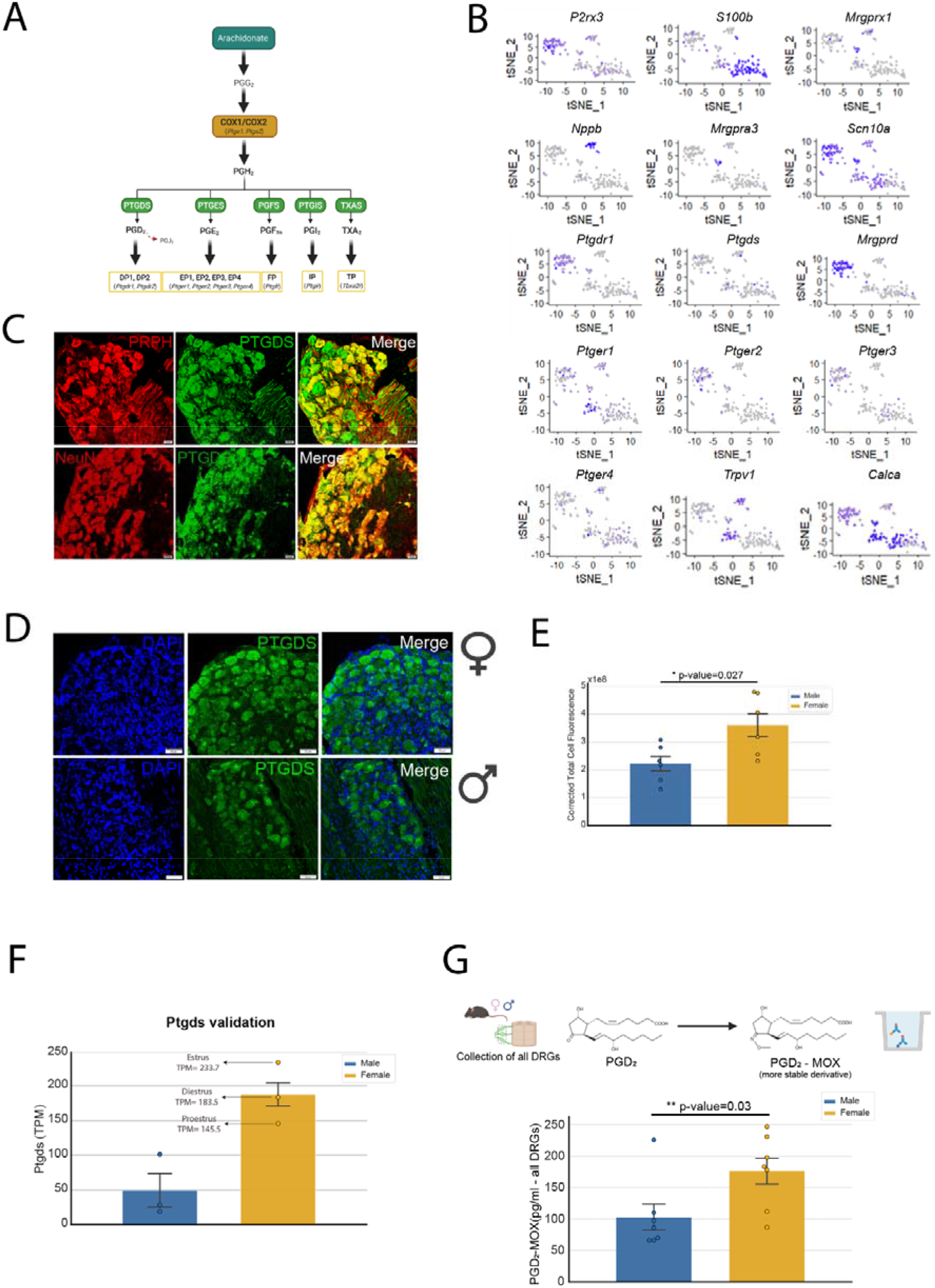
Ptgds expression is higher in female DRGs and leads to higher production of PGD_2_. **A**, Ptgds converts PGH_2_ to PGD_2_, which has a very short half-life and is rapidly metabolized to PGJ_2_. **B**, Single DRG neuron-sequencing shows that *Ptgds* and PGD_2_ receptor DP1 (*Ptgdr1*) are expressed in neurons in the DRG. *Ptgds* is co-expressed with most neuronal markers, suggesting that it is expressed in all neurons; *Ptgdr1* is mostly expressed in non-peptidergic neurons (co-expressed with *P2rx3*); PGE_2_ receptors (*Ptger1*, *Ptger2*, *Ptger3*, *Ptger4*) were expressed by most neuronal subtypes. **C**, We confirmed using IHC that Ptgds is expressed in neurons. **D**, **E**, We found that Ptgds has higher expression in female DRG neurons compared to male DRGs, at the protein level (Unpaired t-test, t = 2.584, df = 10, p-value = 0.0272). **F**, In a separate TRAP experiment, we monitored the female estrous cycle and validated that *Ptgds* is higher in females compared to females regardless of estrous cycle. **G**, PGD_2_-MOX ELISA demonstrated that PGD_2_ levels are higher in female DRGs (Unpaired t-test, t = 2.381, df = 12, p-value = 0.0347). Panel C scale bar = 20 μm; Panel D scale bar = 50 μm.

First, we reanalyzed single-neuron DRG RNA-sequencing (57) and observed *Ptgds* mRNA expression in several subpopulations of neurons, including ones expressing *Calca* (a marker for peptidergic neurons) and *P2rx3* (a marker for non-peptidergic neurons) (**Figure 5B**). We noted that a receptor for PGD_2_, *Ptgdr1* (DP1), was also expressed, especially in non-peptidergic neurons, but *Ptgdr2* (DP2) was not expressed in DRG neurons. We confirmed, using IHC, that Ptgds is expressed in almost all neurons in the mouse DRG (**Figure 5C**). Next, we sought to verify whether there were any sex differences in Ptgds expression at the protein level in the mouse DRG. Ptgds expression was markedly higher in female DRG (**Figure 5D, E**). To verify our previous TRAP experiment, we conducted an independent experiment where we tracked the estrous cycle stage in the female mice. We found that *Ptgds* mRNA associated with ribosomes was substantially higher in this group of female mice with some variation within the estrous cycle, with highest at estrous phase and estrogen levels (**Figure 5F**). In addition, we also investigated whether Ptgds showed sex-differences in expression in brain neurons (**Supp. Figure 1A)**. Similar to observations in the DRG, Ptgds expression was higher in female cortical neurons (**Supp. Figure 1B**).

Having confirmed that Ptgds is more highly expressed in female DRG neurons, we investigated whether this would lead to functional sex differences. First, we measured PGD_2_ levels in female and male DRGs. Because PGD_2_ is highly unstable and can degrade very rapidly, we opted for converting PGD_2_ to a more stable derivative PGD_2_-MOX (by treating our samples with methoxylamine hydrochloride, MOX HCl) and performed an ELISA assay. Our results demonstrated that PGD_2_ levels are higher in female DRGs (**Figure 5G**) in concordance with the higher levels of Ptgds. Next, we tested whether inhibiting Ptgds would produce differential behavioral effects in male and female mice. Unexpectedly, in pilot experiments, we noted intense grimacing behavior in male mice, so we focused on this behavioral output since it is driven by nociceptor input to the CNS (58). We used three different doses (1 mg/kg, 3 mg/kg and 10 mg/kg) of AT-56, a selective and competitive inhibitor of Ptgds (59) observing a robust grimacing effect after intraperitoneal (i.p.) injections of AT-56 (10 mg/kg) in particular in male mice (**Figure 6A**). At each dose, female mice exhibited less grimacing that lasted for a shorter time (**Figure 6B**). This finding suggests that females are protected against Ptgds inhibition-evoked pain because they have higher basal PGD_2_ levels and more enzyme.

**Figure 6:**
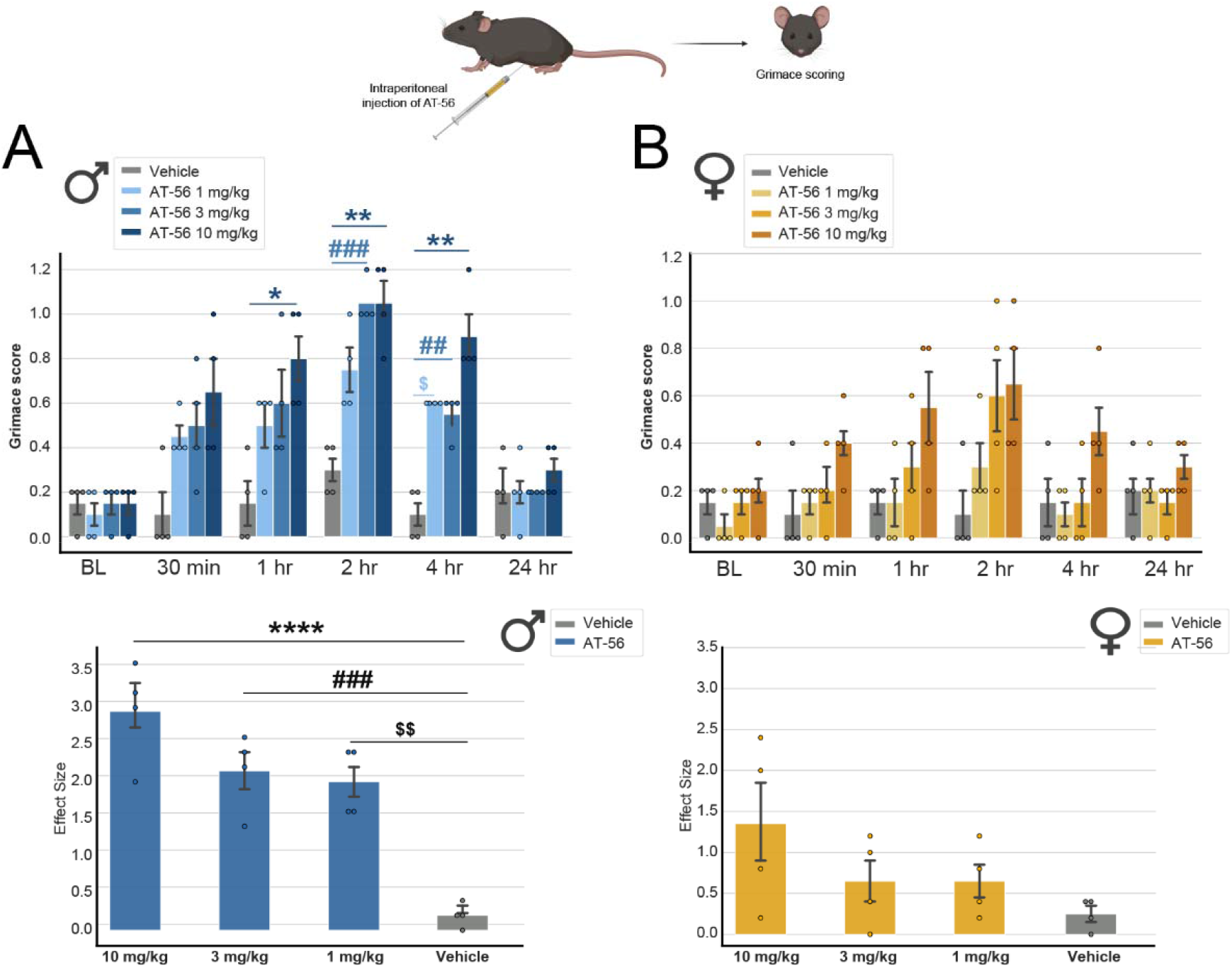
Inhibition of Ptgds produces robust grimacing behavior in mice that is greater in males. **A**, Intraperitoneal injection of AT-56, a selective inhibitor of Ptgds, led to grimacing behavior in male mice (Two-way ANOVA, F = 4.279, p-value<0.0001, post-hoc Sidak’s, * Vehicle vs. AT-56 10 mg/kg male at 1h, p-value= 0.0314; ** Vehicle vs. AT-56 10 mg/kg male at 2h, p-value= 0.0071; *** Vehicle vs. AT-56 10 mg/kg male at 4h, p-value = 0.0068; ### Vehicle vs. AT-56 3 mg/kg male at 2h, p-value = 0.0004; ## Vehicle vs. AT-56 3 mg/kg male at 4h, p-value = 0.0068; $ Vehicle vs. AT-56 1 mg/kg male at 4h, p-value = 0.0193). We also calculated the effect size (difference from the baseline) and observed a significant difference for the 3 doses of AT-56 compared to vehicle in males (One way-ANOVA, F=22.00, p-value<0.0001, post-hoc Tukey’s; **** 10 mg/kg vs. Vehicle, p-value = <0.0001; **** 10 mg/kg vs. Vehicle p-value = <0.0001; ### 3 mg/kg vs. Vehicle p-value = 0.0006; $$ 1 mg/kg vs. Vehicle p-value= 0.0012). **B**, Grimacing behavior in female mice following AT-56 injection was not different from vehicle (Two-way ANOVA, F = 1.136, p-value = 0.3462). When calculating the effect size (difference from the baseline), we did not observe any significant differences between groups in female mice (Two-way ANOVA, F = 2.109, p-value = 0.1524).

A previous study demonstrated that Ptgds gene knockout led to a loss of Prostaglandin E_2_ (PGE_2_) evoked mechanical pain hypersensitivity (43), suggesting an interplay between these closely related molecules (**Figure 5A**) in pain signaling. Moreover, a previous clinical study suggested a sex difference in ibuprofen-induced analgesia in an experimental pain model wherein only males showed analgesia in response to this drug, which lowers PGE_2_ levels (60). We investigated whether there were any differences in mechanical behavior or grimace between males and females in response to PGE_2_. While intraplantar injection of PGE_2_ did not lead to any response to von Frey filaments in male mice at doses of 300 ng or 1 μg (**Figure 7A**), it produced mechanical hypersensitivity in female mice (**Figure 7B**). We did not observe any significant grimacing behavior in male mice (**Figure 7C**) after PGE_2_ injection. Female mice, however, displayed grimacing up to 60 min after PGE_2_ injection (**Figure 7D**).

**Figure 7:**
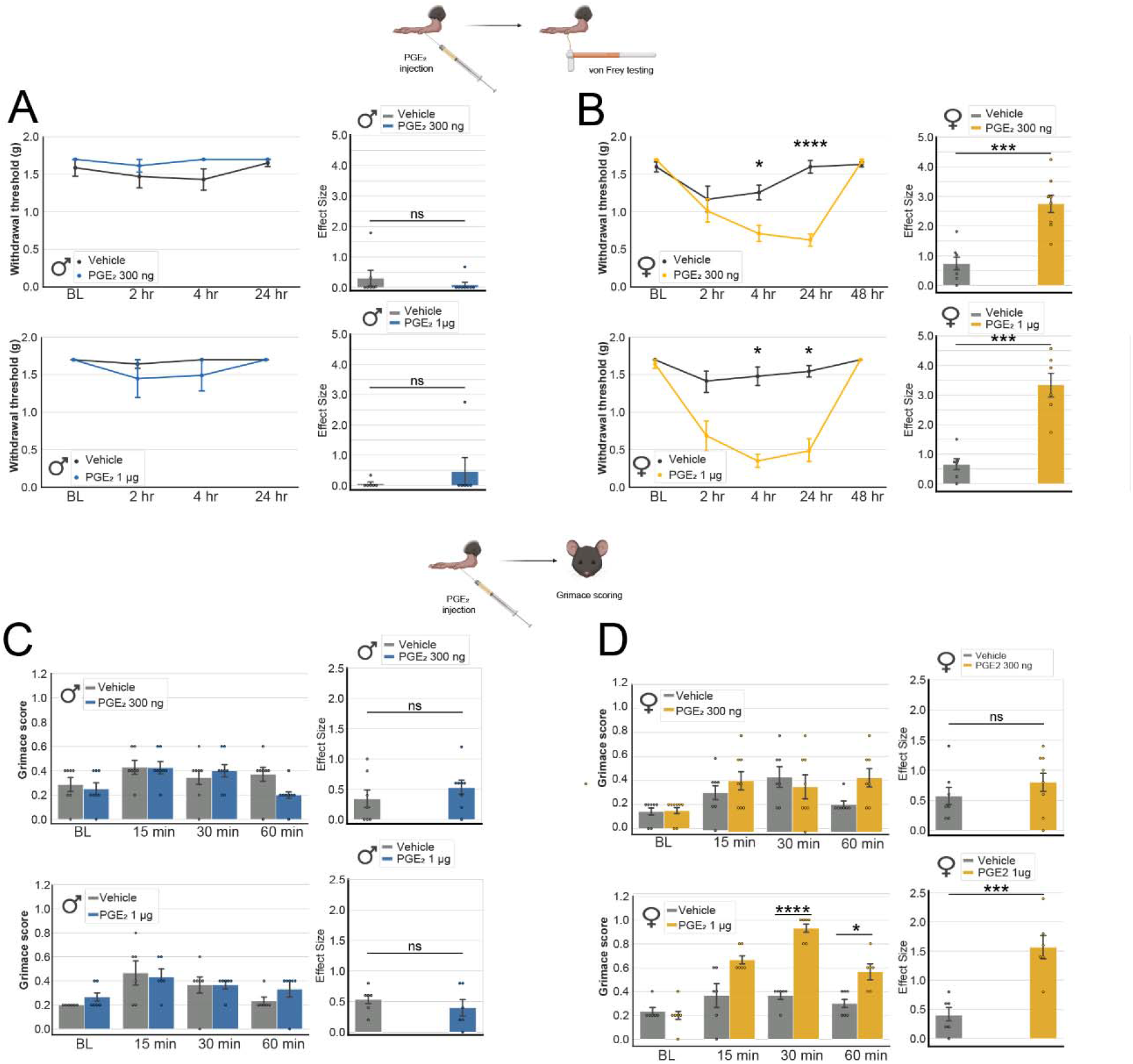
Intraplantar administration of PGE_2_ produces greater mechanical allodynia and grimacing in female mice. **A**, Male mice did not respond to von Frey filaments after injection of 300 ng (Two-way ANOVA RM, F = 1.030, p-value = 0 .3900) or 1 μg (Two-way ANOVA RM, F = 0.7260, p-value = 0.5444) of PGE_2_. When calculating the effect size we did not observe any statistical significant differences between groups in males (Effect size 300 ng: Unpaired t-test, t = 0.8756, df = 13, p-value = 0.3971; Effect size 1 μg: Unpaired t-test, t = 0.8709, df = 10, p-value = 0.4042). **B**, Female mice showed mechanical allodynia up to 24 hours after injection of both 300 ng of PGE_2_ (Two-way ANOVA, F = 12.35, p-value <0.0001, post-hoc Sidak’s: * Vehicle - 300ng PGE_2_ at 4h, p-value = 0.0161, **** Vehicle - 300ng PGE_2_ at 24h, p-value <0.0001) and 1 μg of PGE_2_ (Two-way ANOVA RM, F = 10.78, p-value <0.0001, post-hoc Sidak’s: *** Vehicle - 1 μg PGE_2_ at 4h, p-value = 0.0005, ** Vehicle - 1 μg PGE_2_ at 24h, p-value = 0.0026). We also observed an effect size difference between groups in female mice (*** Effect size 300 ng: Unpaired t-test, t = 4.980, df = 13, p-value = 0.0003; *** Effect size 1 μg: Unpaired t-test, t = 5.569, df = 10, p-value = 0.0002). **C**, Male mice did not show any significant grimacing behaviors following administration of 300 ng of PGE_2_ (Two-way ANOVA RM, F = 1.996, p-value = 0.1305). At 1 μg of PGE_2_ we also did not observe any significant grimacing in male mice (Two-way ANOVA RM, F = 0.5376, p-value = 0.6601). Similarly, we also do not observe any effect size between groups in grimacing of male mice (Effect size 300 ng: Unpaired t-test, t = 0.9225, df = 13, p-value = 0.3731; Effect size 1 μg: Unpaired t-test, t = 0.8305, df = 10, p-value = 0.4257). **D**, Female mice exhibit robust grimacing after 1 μg PGE_2_ injection (Two-way ANOVA RM, F = 11.34, p-value <0.0001, post-hoc Sidak’s: **** Vehicle - 1ug PGE_2_ at 30 min, p-value<0.0001, * Vehicle - 1ug PGE_2_ at 60 min, p-value = 0.0257) but not at 300 ng (Two-way ANOVA RM, F = 1.197, p-value = 0.1303). We also observed an effect size in the grimacing scores between 1 μg PGE_2_ and vehicle grimacing in female mice (Effect size 300 ng: Unpaired t-test, t = 0.9458, df = 13, p-value = 0.3615; *** Effect size 1 μg: Unpaired t-test, t = 4.669, df = 10, p-value = 0.0009).

## DISCUSSION

We used nociceptor TRAP sequencing to reveal sex differences in the translatomes of these neurons that are crucial for nociception. We reach several conclusions based on our work. Consistent with previous studies done at the whole transcriptome level for the DRG, most differences can be attributed to immune genes (36, 38, 61), a finding that may be important for chronic pain(15). At the translatome level, we observed differences in mRNAs bound by ribosomes in nociceptors between males and females, and many of these likely play an important role in the function of these cells. The nociceptor specificity of our TRAP-seq approach, did not identify similar sex differences at the transcriptome and translatome levels for autosomal genes, supporting the important role of translation regulation in nociception (62). We validated higher Ptgds expression in female neurons and found that this leads to a profound sex difference in behavioral response to PGD_2_. We also found sex differences in behavioral responses to PGE_2_ demonstrating that there is a fundamental divergence in prostaglandin signaling between males and females in relation to nociception.

There is a growing body of evidence of mechanistic sex differences in nociception and pain (13, 63, 64). Immune cells have been shown to play an important role in the development and resolution of chronic pain, many of them with sex-specific mechanisms (e.g. macrophages in male mice (65)), where different genes are up-regulated following injury in male and female mice (15). We identified non-neuronal genes with potential roles in inflammatory and immune responses that had differential baseline expression between males and females. These included genes that regulate T cell signaling, such as *Cxcl16* (up-regulated in females) and *Cd276* (up-regulated in males). These findings are consistent with previous literature showing sex differential roles of T cells in opioid analgesia (66) and where distinct proteins produced by T cells are up-regulated following spared nerve injury - PPARα in male mice and PPARγ in female mice (67). Interestingly, in the latter study, it was reported that T cells play a key role in driving neuropathic pain in females (67) whereas monocyte-associated genes such as toll-like receptor 4 (TLR4) contributed to neuropathic pain in males (67, 68).

While sex differences in immune contributions to pain have garnered extensive attention, differences at the neuronal level have not been found in some transcriptomic studies (36), and have not been examined in others (69). However, some recent studies examining specific neuronal populations have found sex differences (16), and some genes, such as the prolactin receptor, show sex differences in expression within neuronal populations (70). Using the TRAP technique, which we have previously used to characterize nociceptor translatome changes in neuropathic pain (22) and between ganglia (23), we have identified a number of sex differences in translation of mRNAs in DRG nociceptors. *Inpp5d* (Inositol polyphosphate-5-phosphatase D) was up-regulated in female nociceptors. This gene has been associated with dementia and Alzheimer’s (71, 72), which are both more common in women (8). It would be of interest to investigate whether translation of this gene is also up-regulated in the brain of females. In contrast, *Apln* (Apelin), which is a neuroprotectant neuropeptide and anti-inflammatory protein, is up-regulated in male nociceptors and it has been reported as a promising target to treat Alzheimer’s disease (73). We also identified several genes with sex differences in mRNA association with ribosomes at the baseline level that have been previously linked to pain (see table 4). For instance, genes in the MAPK cascade have been linked to inflammatory responses in sensory neurons (74), with *Map3k1* (mitogen-activated protein kinase kinase kinase 1) being up-regulated in a model of carrageenan-induced hyperalgesia (75). *Map3k1* was up-regulated in female nociceptors in our study while *Mapk1ip1* (mitogen-activated protein kinase 1 interacting protein 1) was up-regulated in male nociceptors. These findings suggest that although MAPK signalling likely plays a key role in pain in both males and females (76), there may be underlying nuances in signalling that are sex-specific, such as the prominent role of p38 in pain signalling in males (77–79).

We saw little overlap in transcriptome differences in the DRG between males and females and nociceptor translatome differences between males and females. Some of the reason for this is almost certainly technical; however, this divergence might also be partially explained by sex differential translation regulation. This area of research has not garnered much attention but may be important for neuronal function given the key role of translation regulation in synaptic plasticity (80). In the context of pain, we recently demonstrated that female-selective translation of the prolactin receptor mRNA in nociceptors is a causative factor in the pain promoting effects of prolactin in female mice (18).

The increased abundance of *Ptgds* mRNA in the active translatome of female nociceptors intrigued us due to the well-known role of prostaglandins in pain signaling. A previous study showed higher Ptgds expression in the neonatal female brain (81) but we are not aware of any other reports of sex differences for this enzyme in neuronal tissue. Ptgds converts PGH_2_ to PGD_2_, which is one of the most abundant prostaglandins in the brain (41, 42, 82). PGD_2_ is known to have important roles in the regulation of nociception (43, 83), temperature (53) and sleep (52). There is also evidence that Ptgds is involved in the transport of retinoids in the brain (84), thus playing essential roles in the nervous system. In behavioral experiments, inhibition of Ptgds caused robust pain behavior in male mice, while female mice showed effects only at high doses of inhibitor. Higher baseline levels of Ptgds and PGD_2_ in female nociceptors likely explain these sex differences in response to AT-56. Previous studies have shown both anti-(46) (47) and pro-nociceptive roles of PGD_2_ (48). We were not able to test the effect of AT-56 in pain models due to the effect of the drug alone. Interestingly, we also found higher levels of Ptgds protein in female cortex, indicating that our nociceptor findings may be generalizable to other neuronal populations.

Mice lacking the *Ptgds* gene from birth do not develop tactile pain following PGE_2_ injection (43). This suggests an interaction between the *Ptgds* gene and PGE_2_ that prompted us to look for sex differences in response to PGE_2_. We found that female mice responded to lower doses of PGE_2_ with mechanical hypersensitivity and grimacing than did male mice, reminiscent of similar recent findings with calcitonin gene-related peptide (85). This suggests a complex balance between PGD_2_ and PGE_2_ in nociceptive signaling that will take more work to fully understand. Nevertheless, these sex differences in prostaglandin signaling have implications for some of the most commonly used pain relievers – the cyclooxygenase (COX) inhibitors. Studies in rodent arthritis models have described reduced inflammation in COX isoform knockout mice in females but not males (86). In a study using an experimental pain model in healthy human subjects, ibuprofen produced analgesia in men but not women despite equal blood levels of the drug (60). Collectively, these results point to profound sex differences in prostaglandin signaling in the nervous system. Our findings of enhanced translation of *Ptgds* mRNA in the female nervous system can help to understand the molecular underpinnings of these differences better. Nearly a century after the discovery of prostaglandins (87), and centuries after humans started targeting them for pain, mechanistic sex differences in their actions are only beginning to come into focus. In our view, this is a testament to the dire need for consideration of sex as a biological variable in basic neuroscience research.

## REFERENCES

1. Beery AK, Zucker I (2011): Sex bias in neuroscience and biomedical research. Neuroscience & Biobehavioral Reviews. 35:565–572.

2. Mendrek A, Mancini-Marïe A (2016): Sex/gender differences in the brain and cognition in schizophrenia. Neurosci Biobehav Rev. 67:57–78.

3. Nawka A, Kalisova L, Raboch J, Giacco D, Cihal L, Onchev G, et al. (2013): Gender differences in coerced patients with schizophrenia. BMC Psychiatry. 13:257–257.

4. Baldereschi M, Di Carlo A, Rocca WA, Vanni P, Maggi S, Perissinotto E, et al. (2000): Parkinson’s disease and parkinsonism in a longitudinal study: two-fold higher incidence in men. ILSA Working Group. Italian Longitudinal Study on Aging. Neurology. 55:1358–1363.

5. Georgiev D, Hamberg K, Hariz M, Forsgren L, Hariz GM (2017): Gender differences in Parkinson’s disease: A clinical perspective. Acta Neurol Scand. 136:570–584.

6. Accortt EE, Freeman MP, Allen JJ (2008): Women and major depressive disorder: clinical perspectives on causal pathways. J Womens Health (Larchmt). 17:1583–1590.

7. Bruce SE, Yonkers KA, Otto MW, Eisen JL, Weisberg RB, Pagano M, et al. (2005): Influence of psychiatric comorbidity on recovery and recurrence in generalized anxiety disorder, social phobia, and panic disorder: a 12-year prospective study. Am J Psychiatry. 162:1179–1187.

8. Gao S, Burney HN, Callahan CM, Purnell CE, Hendrie HC (2019): Incidence of Dementia and Alzheimer Disease Over Time: A Meta-Analysis. J Am Geriatr Soc. 67:1361–1369.

9. Mogil JS (2012): Sex differences in pain and pain inhibition: multiple explanations of a controversial phenomenon. Nat Rev Neurosci. 13:859–866.

10. Fillingim RB, King CD, Ribeiro-Dasilva MC, Rahim-Williams B, Riley JL, 3rd (2009): Sex, gender, and pain: a review of recent clinical and experimental findings. J Pain. 10:447–485.

11. Arout CA, Sofuoglu M, Bastian LA, Rosenheck RA (2018): Gender Differences in the Prevalence of Fibromyalgia and in Concomitant Medical and Psychiatric Disorders: A National Veterans Health Administration Study. Journal of women’s health (2002). 27:1035–1044.

12. Torrance N, Smith BH, Bennett MI, Lee AJ (2006): The epidemiology of chronic pain of predominantly neuropathic origin. Results from a general population survey. The Journal of Pain. 7:281–289.

13. Ruau D, Liu LY, Clark JD, Angst MS, Butte AJ (2012): Sex differences in reported pain across 11,000 patients captured in electronic medical records. J Pain. 13:228–234.

14. Shansky RM, Woolley CS (2016): Considering Sex as a Biological Variable Will Be Valuable for Neuroscience Research. J Neurosci. 36:11817–11822.

15. Mogil JS (2020): Qualitative sex differences in pain processing: emerging evidence of a biased literature. Nat Rev Neurosci. 21:353–365.

16. Smith-Anttila CJA, Mason EA, Wells CA, Aronow BJ, Osborne PB, Keast JR (2020): Identification of a Sacral, Visceral Sensory Transcriptome in Embryonic and Adult Mice. eNeuro. 7.

17. Liu Y, Beyer A, Aebersold R (2016): On the Dependency of Cellular Protein Levels on mRNA Abundance. Cell. 165:535–550.

18. Patil M, Belugin S, Mecklenburg J, Wangzhou A, Paige C, Barba-Escobedo PA, et al. (2019): Prolactin Regulates Pain Responses via a Female-Selective Nociceptor-Specific Mechanism. iScience. 20:449–465.

19. Patil MJ, Green DP, Henry MA, Akopian AN (2013): Sex-dependent roles of prolactin and prolactin receptor in postoperative pain and hyperalgesia in mice. Neuroscience. 253:132–141.

20. Patil MJ, Ruparel SB, Henry MA, Akopian AN (2013): Prolactin regulates TRPV1, TRPA1, and TRPM8 in sensory neurons in a sex-dependent manner: Contribution of prolactin receptor to inflammatory pain. Am J Physiol Endocrinol Metab. 305:E1154–1164.

21. Heiman M, Schaefer A, Gong S, Peterson JD, Day M, Ramsey KE, et al. (2008): A translational profiling approach for the molecular characterization of CNS cell types. Cell. 135:738–748.

22. Megat S, Ray PR, Moy JK, Lou TF, Barragan-Iglesias P, Li Y, et al. (2019): Nociceptor Translational Profiling Reveals the Ragulator-Rag GTPase Complex as a Critical Generator of Neuropathic Pain. J Neurosci. 39:393–411.

23. Megat S, Ray PR, Tavares-Ferreira D, Moy JK, Sankaranarayanan I, Wanghzou A, et al. (2019): Differences between Dorsal Root and Trigeminal Ganglion Nociceptors in Mice Revealed by Translational Profiling. J Neurosci. 39:6829–6847.

24. Wilcox CM, Cryer B, Triadafilopoulos G (2005): Patterns of use and public perception of over-the-counter pain relievers: focus on nonsteroidal antiinflammatory drugs. J Rheumatol. 32:2218–2224.

25. Dobin A, Davis CA, Schlesinger F, Drenkow J, Zaleski C, Jha S, et al. (2013): STAR: ultrafast universal RNA-seq aligner. Bioinformatics. 29:15–21.

26. Pertea M, Pertea GM, Antonescu CM, Chang T-C, Mendell JT, Salzberg SL (2015): StringTie enables improved reconstruction of a transcriptome from RNA-seq reads. Nature biotechnology. 33:290–295.

27. Bhattacharyya, A (1943): On a measure of divergence between two statistical populations defined by their probability distributions. Bull Calcutta math Soc. 35:99–109.

28. Zhang XD (2011): Illustration of SSMD, z score, SSMD*, z* score, and t statistic for hit selection in RNAi high-throughput screens. Journal of Biomolecular Screening. 16:775–785.

29. Zhang XD, Lacson R, Yang R, Marine SD, McCampbell A, Toolan DM, et al. (2010): The Use of SSMD-Based False Discovery and False Nondiscovery Rates in Genome-Scale RNAi Screens. Journal of Biomolecular Screening. 15:1123–1131.

30. Zhou P, Zhang Y, Ma Q, Gu F, Day DS, He A, et al. (2013): Interrogating translational efficiency and lineage-specific transcriptomes using ribosome affinity purification. Proceedings of the National Academy of Sciences. 110:15395–15400.

31. Megat S, Ray PR, Moy JK, Lou T-F, Barragan-Iglesias P, Li Y, et al. (2019): Nociceptor translational profiling reveals the ragulator-Rag GTPase complex as a critical generator of neuropathic pain. Journal of Neuroscience. 39:393–411.

32. Ray P, Torck A, Quigley L, Wangzhou A, Neiman M, Rao C, et al. (2018): Comparative transcriptome profiling of the human and mouse dorsal root ganglia: an RNA-seq-based resource for pain and sensory neuroscience research. Pain. 159:1325.

33. Obata K, Yamanaka H, Fukuoka T, Yi D, Tokunaga A, Hashimoto N, et al. (2003): Contribution of injured and uninjured dorsal root ganglion neurons to pain behavior and the changes in gene expression following chronic constriction injury of the sciatic nerve in rats. Pain. 101:65–77.

34. Nguyen MQ, Le Pichon CE, Ryba N (2019): Stereotyped transcriptomic transformation of somatosensory neurons in response to injury. Elife. 8.

35. Mi H, Muruganujan A, Ebert D, Huang X, Thomas PD (2019): PANTHER version 14: more genomes, a new PANTHER GO-slim and improvements in enrichment analysis tools. Nucleic acids research. 47:D419–D426.

36. Lopes DM, Malek N, Edye M, Jager SB, McMurray S, McMahon SB, et al. (2017): Sex differences in peripheral not central immune responses to pain-inducing injury. Sci Rep. 7:16460.

37. Liang Z, Hore Z, Harley P, Stanley FU, Michrowska A, Dahiya M, et al. (2020): A transcriptional toolbox for exploring peripheral neuro-immune interactions. Pain.

38. North RY, Li Y, Ray P, Rhines LD, Tatsui CE, Rao G, et al. (2019): Electrophysiological and transcriptomic correlates of neuropathic pain in human dorsal root ganglion neurons. Brain. 142:1215–1226.

39. Thoreen CC, Chantranupong L, Keys HR, Wang T, Gray NS, Sabatini DM (2012): A unifying model for mTORC1-mediated regulation of mRNA translation. Nature. 485:109–113.

40. Jensen KB, Dredge BK, Toubia J, Jin X, Iadevaia V, Goodall GJ, et al. (2020): capCLIP: a new tool to probe protein synthesis in human cells through capture and identification of the eIF4E-mRNA interactome. bioRxiv.2020.2004.2018.047571.

41. Shaik JS, Miller TM, Graham SH, Manole MD, Poloyac SM (2014): Rapid and simultaneous quantitation of prostanoids by UPLC-MS/MS in rat brain. J Chromatogr B Analyt Technol Biomed Life Sci. 945-946:207–216.

42. Ogorochi T, Narumiya S, Mizuno N, Yamashita K, Miyazaki H, Hayaishi O (1984): Regional distribution of prostaglandins D2, E2, and F2α and related enzymes in postmortem human brain. Journal of neurochemistry. 43:71–82.

43. Eguchi N, Minami T, Shirafuji N, Kanaoka Y, Tanaka T, Nagata A, et al. (1999): Lack of tactile pain (allodynia) in lipocalin-type prostaglandin D synthase-deficient mice. Proc Natl Acad Sci U S A. 96:726–730.

44. Ito S, Okuda-Ashitaka E, Minami T (2001): Central and peripheral roles of prostaglandins in pain and their interactions with novel neuropeptides nociceptin and nocistatin. Neuroscience research. 41:299–332.

45. Liang X, Wu L, Hand T, Andreasson K (2005): Prostaglandin D2 mediates neuronal protection via the DP1 receptor. J Neurochem. 92:477–486.

46. Minami T, Okuda-Ashitaka E, Mori H, Ito S, Hayaishi O (1996): Prostaglandin D2 inhibits prostaglandin E2-induced allodynia in conscious mice. Journal of Pharmacology and Experimental Therapeutics. 278:1146–1152.

47. Minami T, Okuda-Ashitaka E, Nishizawa M, Mori H, Ito S (1997): Inhibition of nociceptin-induced allodynia in conscious mice by prostaglandin D2. British journal of pharmacology. 122:605.

48. Ohkubo T, Shibata M, Takahashi H, Inoki R (1983): Effect of prostaglandin D2 on pain and inflammation. Jpn J Pharmacol. 33:264–266.

49. Sekeroglu A, Jacobsen JM, Jansen-Olesen I, Gupta S, Sheykhzade M, Olesen J, et al. (2017): Effect of PGD(2) on middle meningeal artery and mRNA expression profile of L-PGD(2) synthase and DP receptors in trigeminovascular system and other pain processing structures in rat brain. Pharmacol Rep. 69:50–56.

50. Shimizu T, Mizuno N, Amano T, Hayaishi O (1979): Prostaglandin D2, a neuromodulator. Proceedings of the National Academy of Sciences. 76:6231–6234.

51. Uda R, Horiguchi S, Ito S, Hyodo M, Hayaishi O (1990): Nociceptive effects induced by intrathecal administration of prostaglandin D2, E2, or F2 alpha to conscious mice. Brain Res. 510:26–32.

52. Urade Y, Hayaishi O (1999): Prostaglandin D2 and sleep regulation. Biochimica et Biophysica Acta (BBA)-Molecular and Cell Biology of Lipids. 1436:606–615.

53. Wang TA, Teo CF, Åkerblom M, Chen C, Tynan-La Fontaine M, Greiner VJ, et al. (2019): Thermoregulation via temperature-dependent PGD2 production in mouse preoptic area. Neuron. 103:309–322. e307.

54. Ferrer MD, Busquets-Cortés C, Capó X, Tejada S, Tur JA, Pons A, et al. (2019): Cyclooxygenase-2 Inhibitors as a Therapeutic Target in Inflammatory Diseases. Curr Med Chem. 26:3225–3241.

55. Sala A, Proschak E, Steinhilber D, Rovati GE (2018): Two-pronged approach to anti-inflammatory therapy through the modulation of the arachidonic acid cascade. Biochem Pharmacol. 158:161–173.

56. Marjoribanks J, Ayeleke RO, Farquhar C, Proctor M (2015): Nonsteroidal anti-inflammatory drugs for dysmenorrhoea. Cochrane Database Syst Rev. 2015:Cd001751.

57. Hu G, Huang K, Hu Y, Du G, Xue Z, Zhu X, et al. (2016): Single-cell RNA-seq reveals distinct injury responses in different types of DRG sensory neurons. Sci Rep. 6:31851.

58. Langford DJ, Bailey AL, Chanda ML, Clarke SE, Drummond TE, Echols S, et al. (2010): Coding of facial expressions of pain in the laboratory mouse. Nat Methods. 7:447–449.

59. Irikura D, Aritake K, Nagata N, Maruyama T, Shimamoto S, Urade Y (2009): Biochemical, functional, and pharmacological characterization of AT-56, an orally active and selective inhibitor of lipocalin-type prostaglandin D synthase. Journal of Biological Chemistry. 284:7623–7630.

60. Walker JS, Carmody JJ (1998): Experimental pain in healthy human subjects: gender differences in nociception and in response to ibuprofen. Anesth Analg. 86:1257–1262.

61. Lopes DM, Denk F, McMahon SB (2017): The Molecular Fingerprint of Dorsal Root and Trigeminal Ganglion Neurons. Front Mol Neurosci. 10:304.

62. Khoutorsky A, Price TJ (2018): Translational Control Mechanisms in Persistent Pain. Trends Neurosci. 41:100–114.

63. Fillingim RB (2017): Sex, Gender, and Pain. Elsevier, pp 481–496.

64. Mogil JS (2020): Qualitative sex differences in pain processing: emerging evidence of a biased literature. Nature Reviews Neuroscience. 21:353–365.

65. Yu X, Liu H, Hamel KA, Morvan MG, Yu S, Leff J, et al. (2020): Dorsal root ganglion macrophages contribute to both the initiation and persistence of neuropathic pain. Nat Commun. 11:264.

66. Rosen SF, Ham B, Haichin M, Walters IC, Tohyama S, Sotocinal SG, et al. (2019): Increased pain sensitivity and decreased opioid analgesia in T-cell-deficient mice and implications for sex differences. Pain. 160:358–366.

67. Sorge RE, Mapplebeck JC, Rosen S, Beggs S, Taves S, Alexander JK, et al. (2015): Different immune cells mediate mechanical pain hypersensitivity in male and female mice. Nat Neurosci. 18:1081–1083.

68. Sorge RE, LaCroix-Fralish ML, Tuttle AH, Sotocinal SG, Austin JS, Ritchie J, et al. (2011): Spinal cord Toll-like receptor 4 mediates inflammatory and neuropathic hypersensitivity in male but not female mice. J Neurosci. 31:15450–15454.

69. Zeisel A, Hochgerner H, Lonnerberg P, Johnsson A, Memic F, van der Zwan J, et al. (2018): Molecular Architecture of the Mouse Nervous System. Cell. 174:999–1014.e1022.

70. Patil M, Hovhannisyan AH, Wangzhou A, Mecklenburg J, Koek W, Goffin V, et al. (2019): Prolactin receptor expression in mouse dorsal root ganglia neuronal subtypes is sex-dependent. J Neuroendocrinol. 31:e12759.

71. Kim JH (2018): Genetics of Alzheimer’s Disease. Dement Neurocogn Disord. 17:131–136.

72. Yoshino Y, Yamazaki K, Ozaki Y, Sao T, Yoshida T, Mori T, et al. (2017): INPP5D mRNA Expression and Cognitive Decline in Japanese Alzheimer’s Disease Subjects. J Alzheimers Dis. 58:687–694.

73. Masoumi J, Abbasloui M, Parvan R, Mohammadnejad D, Pavon-Djavid G, Barzegari A, et al. (2018): Apelin, a promising target for Alzheimer disease prevention and treatment. Neuropeptides. 70:76–86.

74. Ji RR (2004): Peripheral and central mechanisms of inflammatory pain, with emphasis on MAP kinases. Curr Drug Targets Inflamm Allergy. 3:299–303.

75. Yukhananov R, Kissin I (2008): Persistent changes in spinal cord gene expression after recovery from inflammatory hyperalgesia: a preliminary study on pain memory. BMC Neurosci. 9:32–32.

76. Ji RR, Gereau RWt, Malcangio M, Strichartz GR (2009): MAP kinase and pain. Brain Res Rev. 60:135–148.

77. Taves S, Berta T, Liu DL, Gan S, Chen G, Kim YH, et al. (2016): Spinal inhibition of p38 MAP kinase reduces inflammatory and neuropathic pain in male but not female mice: Sex-dependent microglial signaling in the spinal cord. Brain Behav Immun. 55:70–81.

78. Mapplebeck JCS, Dalgarno R, Tu Y, Moriarty O, Beggs S, Kwok CHT, et al. (2018): Microglial P2X4R-evoked pain hypersensitivity is sexually dimorphic in rats. Pain. 159:1752–1763.

79. Paige C, Maruthy GB, Mejia G, Dussor G, Price T (2018): Spinal Inhibition of P2XR or p38 Signaling Disrupts Hyperalgesic Priming in Male, but not Female, Mice. Neuroscience. 385:133–142.

80. Biever A, Donlin-Asp PG, Schuman EM (2019): Local translation in neuronal processes. Curr Opin Neurobiol. 57:141–148.

81. Sakakibara M, Uenoyama Y, Minabe S, Watanabe Y, Deura C, Nakamura S, et al. (2013): Microarray analysis of perinatal-estrogen-induced changes in gene expression related to brain sexual differentiation in mice. PloS one. 8:e79437.

82. Hertting G, Seregi A (1989): Formation and function of eicosanoids in the central nervous system. Ann N Y Acad Sci. 559:84–99.

83. Bellei E, Monari E, Bergamini S, Cuoghi A, Tomasi A, Guerzoni S, et al. (2015): Validation of potential candidate biomarkers of drug-induced nephrotoxicity and allodynia in medication-overuse headache. J Headache Pain. 16:559–559.

84. Tanaka T, Urade Y, Kimura H, Eguchi N, Nishikawa A, Hayaishi O (1997): Lipocalin-type prostaglandin D synthase (beta-trace) is a newly recognized type of retinoid transporter. J Biol Chem. 272:15789–15795.

85. Avona A, Burgos-Vega C, Burton MD, Akopian AN, Price TJ, Dussor G (2019): Dural Calcitonin Gene-Related Peptide Produces Female-Specific Responses in Rodent Migraine Models. J Neurosci. 39:4323–4331.

86. Chillingworth NL, Morham SG, Donaldson LF (2006): Sex differences in inflammation and inflammatory pain in cyclooxygenase-deficient mice. Am J Physiol Regul Integr Comp Physiol. 291:R327–334.

87. v. Euler US (1935): Über die Spezifische Blutdrucksenkende Substanz des Menschlichen Prostata-und Samenblasensekretes. Klinische Wochenschrift. 14:1182–1183.

