## Supplemental Information for "Sex differences in nociceptor translatomes contribute to divergent prostaglandin signaling in male and female mice"

**Supplemental Materials**

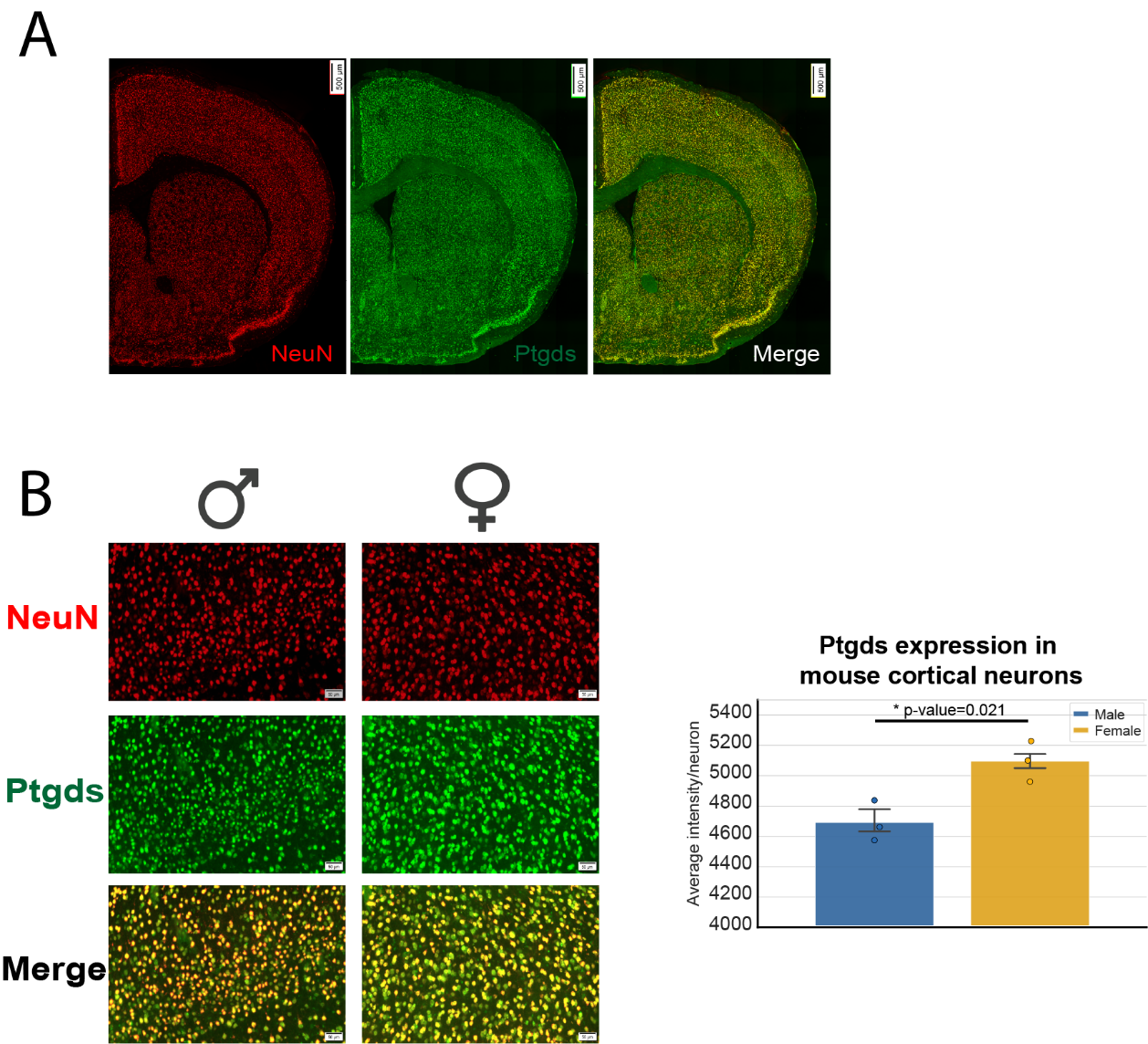

**Suppl Figure 1:** *Ptgds expression is higher in female cortical neurons.* **A**, Ptgds is highly co-localized with neuronal marker NeuN in the mouse brain. **B**, Ptgds expression in mouse cortical neurons is higher in females compared to males (* Unpaired t-test, t = 3.699, df = 4, p-value= 0.0209). A total of 100 neurons per slice (3 slices per animal) were used to quantify Ptgds. Panel A Scale bar = 500 μm; panel B scale bar = 50 μm.

| **Gene name** | **Gene description** | **GO Biological Process** | **GO Cellular Component** | **GO Molecular Function** | **Nervous system; nociception related?** | **Reference** |
| --- | --- | --- | --- | --- | --- | --- |
|  | **UP-REGULATED IN FEMALES (INPUT)** | | | | | |
| **Hoxd4** | Homeobox d4 | Transcription regulation | Nuceloplasm; Cell junction | Transcription factor | Wallerian degeneration | PMID: 31397358 |
| **Smpd4** | Sphingomyelin phosphodiesterase 4 | Sphingomyelin catabolic process | Nucleus; Membrane; Golgi apparatus; Endoplasmic reticulum | Sphingomyelin phosphodiesterase activity | Loss of this protein is associated with microcephaly | PMID: 31495489 |
| **Vpreb3** | Pre-b lymphocyte gene 3 | Regulation of immunoglobulin secretion | Endoplasmic reticulum; extracellular space | *ND* | *ND* |  |
| **Bc051142** | cDNA sequence bc051142 | *ND* | Membrane | *ND* |  |  |
| **Runx1t1** | Runx1 translocation partner 1 | Transcription regulation | Nucleus | Transcription factor activity | Neuronal differentiation in hippocampus | PMID: 25473084 |
| **Adra2b** | Adrenergic receptor, alpha 2b | MAPK casacade; Regulation of vasoconstriction; Regulation of muscle contraction and neuronal differentiation | Membrane | Adrenergic receptor activity; epinephrine binding | *ND* |  |
| **Srp19** | Signal recognition particle 19 | Srp-dependent cotranslational protein targeting to membrane | Nucleus; Signal recognition particle; Endoplasmic reticulum | 7s RNA binding; ribosome binding | *ND* |  |
| **Lepr** | Leptin receptor | Regulation of mapk cascade; signal transduction; protein phosphorylation | Membrane | Transmembrane signaling receptor activity | Neuropathic pain | PMC5651484; PMID: 28822786 |
| **Kng2** | Kininogen 2 | Regulation of endopeptidase activity; Regulation of cytosolic calcium concentration | Extracellular space | Endopeptidase inhibitor activity; Bradykinin precursor | *ND* |  |
| **Chrm5** | Cholinergic receptor, muscarinic 5 | Transmission of nerve impulse; Chemical synaptic transmission | Synapse; Membrane; Cell junction | G protein-coupled acetylcholine receptor activity; Extracellularly glycine-gated chloride channel activity | Opioid- or nausea/vomiting signaling pathways | PMID: 21570824 |
| **Anpep** | Alanyl (membrane) aminopeptidase | Angiogenesis; Cell differentiation | Membrane; Cytoplasm; Extracellular space | Aminopeptidase activity | Complex regional pain syndrome | PMID: 24244504 |
| **Tmem230** | Transmembrane protein 230 | Synaptic vesicle transport | Membrane; Synaptic vesicle; autophagosome | *ND* | Parkinson’s disease | PMID: 30175983 |
| **Scube3** | Signal peptide, cub domain, egf-like 3 | Regulation of smoothened signaling pathway | Membrane; Extracellular space | Calcium ion binding; protein binding |  |  |
| **Plcb4** | Phospholipase c, beta 4 | Modulation of chemical synaptic transmission; metabolic process | Cell; Nucleus; Cytoplasm; Postsynapse | Hydrolase activity; protein binding | Inflammatory pain | PMID: 12954872 |
| **Kif23** | Kinesin family member 23 | regulation of cytokinesis | Cell; Nucleoplasm | Microtubule binding; ATP binding | *ND* |  |
| **Bc049762** | cDNA sequence Bc049762 | *ND* | *ND* | *ND* | *ND* |  |
| **Ackr3** | Atypical chemokine receptor 3 | Cell adhesion; Immune response | Cell; Cytoplasm; Membrane; Nucleus | Scavenger receptor activity; Chemokine receptor activity | *ND* |  |
| **Sirpb1b** | Signal-regulatory protein beta 1b | Cell-cell adhesion; phagocytosis; T cell activation | Membrane | *ND* | *ND* |  |
| **Pamr1** | Peptidase domain containing associated with muscle regeneration 1 | Proteolysis | Extracellular space | Endopeptidase activity; calcium ion binding | *ND* |  |
| **Jchain** | Immunoglobulin joining chain | Immune response | Extracellular space | Immunogloblobulin binding | *ND* |  |
| **Pi16** | Peptidase inhibitor 16 | Peptidase activity | Extracellular space | Peptidase inhibitor activity | Fibroblast-derived protein involved in neuropathic pain | PMID: 32079726 |
| **Ncoa5** | Nuclear receptor coactivator 5 | Glucose homeostasis | Nucleoplasm; Nucleus | Chromatin binding | *ND* |  |
| **Snurf** | SNRPN upstream reading frame | *ND* | Nucleus | ATPase binding | *ND* |  |
| **Trabd2b** | Trab domain containing 2b | Wnt signaling pathway | Membrane | Hydrolase activity; Peptidase activity | *ND* |  |
| **Card6** | Caspase recruitment domain family, member 6 | I-KappaB kinase/nf-kappaB signaling; Apoptotic process | Cytoplasm | *ND* | Spinal cord injury; Reumathoid arthritis | PMID: 31841440; PMID: 32317187 |
| **Lbp** | Lipopolysaccharide binding protein | Immune response | Membrane; Extracellular space | Lipopolysaccharide binding | *ND* |  |
| **Epb41l5** | Erythrocyte membrane protein band 4.1 like 5 | Epithelial cell morphogenesis | Membrane; Cytoplasm; Cell junction | Cytoskeletal protein binding | *ND* |  |
| **Map3k8** | Mitogen-activated protein kinase kinase kinase 8 | MAPK cascade; Immune system process | Cytoplasm | Kinase activity; Nucleotide binding | *ND* |  |
| **Dio2** | Deiodinase, iodothyronine, type ii | Regulation of cold-induced thermogenesis; Hormone biosynthetic process | Membrane | Oxidoreductase activity | Osteoarthritis | PMID: 25693039 |
| **Nudt6** | Nudix (nucleoside diphosphate linked moiety x)-type motif 6 | Regulation of cell cycle | Cytoplasm | Hydrolase activity | *ND* |  |
| **Pkd2l2** | Polycystic kidney disease 2-like 2 (also known as trpp5) | Ion transmembrane transport | Membrane | Calcium channel activity | *ND* |  |
| **Kdm5c** | Lysine (k)-specific demethylase 5c | Transcription regulation; chromatin organization | Nucleus; cytoplasm | Transcription factor activity;histone demethylase activity | *ND* |  |
| **Pnoc** | Prepronociceptin | Sensory preception | Neuronal cell body | Opioid peptide activity |  |  |
| **Eif2s3x** | Eukaryotic translation initiation factor 2, subunit 3, structural gene x-linked | Formation of translation preinitiation complex | Eukaryotic translation initiation factor 2 complex in cytoplasm | Translation factor activity; RNA binding | *ND* |  |
| **Catip** | Ciliogenesis associated ttc17 interacting protein | Cell projection organization | Cytoskeleton; membrane; cytoplasm; nucleus | *ND* | *ND* |  |
| **Arg1** | Arginase | Immune response; metabolic process | Neuron projection; cytoplasm; extracellular space | Arginase activity | *ND* |  |
| **Cfap70** | Cilia and flagella associated protein 70 | Cillium movement and assembly | Axoneme | Protein binding | *ND* |  |
| **Tnfrsf11b** | Tumor necrosis factor receptor superfamily, member 11b (osteoprotegerin) | Signal transduction; Extracellular matrix organization | Extracellular space | Protein binding | *ND* |  |
| **Dapl1** | Death associated protein-like 1 | Apoptotic signaling pathway | *ND* | Death domain binding | *ND* |  |
| **Cxcl16** | Chemokine (c-x-c motif) ligand 16 | T cell chemotaxis | Extracellular space; Membrane | Chemokine activity | Inflammatory pain | PMID: 22815815 |
| **Fam221a** | Family with sequence similarity 221, member A | *ND* | *ND* | *ND* | *ND* |  |
|  | **UP-REGULATED IN MALES (INPUT)** | | | | | |
| **Qrfpr** | Pyroglutamylated rfamide peptide receptor | Neuropeptide signaling pathway; Metabolism and energy homeostasis | Membrane | Neuropeptide Y receptor activity | *ND* |  |
| **Spry1** | Sprouty RTK signaling antagonist 1 | MAPK activity; Regulation of fibroblast growth factor receptor signaling pathway | Cytoplasm; Nucleus; Membrane | Protein binding | *ND* |  |
| **BC030867 (Hrob )** | Homologous recombination factor with ob-fold | Cellular response to DNA damage stimulus | Site of DNA damage | Single-stranded DNA binding | *ND* |  |
| **Pappa2** | Pappalysin 2 | Proteolysis | Apical plasma membrane; cytosol | Metallopeptidase activity | *ND* |  |
| **Lctl** | Lactase-like | Response to stimuli; Metabolic process | *ND* | *ND* | *ND* |  |
| **Col10a1** | Collagen, type x, alpha 1 | Extracellular matrix organization | Extracellular space | Extracellular matrix structural constituent | Osteoarthritis | PMID: 20498197 |
| **Pclaf** | Pcna clamp associated factor | Cellular response to DNA damage stimulus | Centrosome; Nucleus | Chromatin binding | *ND* |  |
| **Gp9** | Glycoprotein 9 (platelet) | Cell adhesion; Homeostasis | Membrane | Protein binding | *ND* |  |
| **Foxd3** | Forkhead box d3 | Transcription regulation | Nucleus | Transcription factor activity | *ND* |  |
| **Mrap2** | Melanocortin 2 receptor accessory protein 2 | Energy homeostasis; Metabolic process | Membrane | Receptor regulator activity | *ND* |  |
| **Sfi1** | Sfi1 homolog, spindle assembly associated | Regulation of phosphatase activity | Cytoplasm; Centrosome; Cytoskeleton | Phosphatase binding | *ND* |  |
| **Nmbr** | Neuromedin B receptor | Neuropeptide signaling pathway | Membrane; Cytosol | Neuropeptide receptor activity | Itch transmission; Thermal stimuli perception | PMID: 29133874; PMID: 22723708 |
| **Jmjd8** | Jumonji domain containing 8 | Regulation of I-kappaB kinase/nf-kappaB signaling; Regulation of glycolytic process; Regulation of sprouting angiogenesis | Cytoplasm; Nucleus | *ND* | *ND* |  |
| **Dut** | Deoxyuridine triphosphatase | Regulation of signaling receptor activity; Regulation of protein-containing complex assembly | Cytoplasm; Nucleus; Mithocondrion | Receptor inhibitor activity | *ND* |  |
| **Cd276** | Cd276 antigen | Regulation of inflammatory response; Regulation of cytokine production; T cell receptor signaling pathway | Membrane | Signaling receptor binding | *ND* |  |
| **Srd5a2** | Steroid 5 alpha-reductase 2 | Sex differentiation; Testosterone biosynthetic process | Cytoplasm; Membrane | Testosterone biosynthetic process | *ND* |  |
| **Nat14** | N-acetyltransferase 14 | *ND* | Membrane | Transferase activity | *ND* |  |
| **Crtac1** | Cartilage acidic protein 1 | Axonal fasciculation; Regulation of receptor binding | Extracellular region | Calcium ion binding; Protein binding | Osteoarthritis | PMID: 27415616 |
| **Smtnl2** | Smoothelin-like 2 | Actin cytoskeleton organization | Microtubule organizing center | Protein binding |  |  |
| **Nrarp** | Notch-regulated ankyrin repeat protein | Notch signaling pathway; Regulation of T cell differentiation; Regulation of canonical Wnt signaling pathway; Regulation of cell-cell adhesion | *ND* | Protein binding | *ND* |  |
| **Cdca8** | Cell division cycle associated 8 | Cell cycle and division | Cell | Protein binding |  |  |
| **Ep400** | E1a binding protein p400 | Chromatin organization | Nucleus | Nucleotide binding; Helicase activity | Role in myelination in CNS | PMID: 31081019 |
| **Fgf2** | Fibroblast growth factor 2 | Cell differentiation; regulation of fibroblast migration; angiogenesis | Extracellular space; cell; nucleus | Chemoattractant activity; growth factor activity | Bone cancer pain | PMID: 27506449 |
| **Fscn1** | Fascin actin-bundling protein 1 | Regulation of extracellular matrix; Regulation of extracellular matrix | Cytoplasm; cell-cell junction; Myelin sheath | Protein binding; actin binding | Inflammatory component in neuropathic pain | PMID: 29052088 |
| **Ky** | Kyphoscoliosis peptidase | *ND* | *ND* | *ND* |  |  |
| **Chd7** | Chromodomain helicase DNA binding protein 7 | Central nervous system development; Chromatin organization | Nucleus | Protein binding; Helicase activity; Hydrolase activity | Neuron differentiation; CNS myelination | PMID: 28317875; PMID: 26928066 |
| **Atf3** | Activating transcription factor 3 | Transcription regulation | Nucleus | Transcription factor activity | Injury marker | PMID: 15066140 |
| **Grid2ip** | Glutamate receptor, ionotropic, delta 2 (grid2) interacting protein 1 | G protein-coupled glutamate receptor signaling pathway; Actin cytoskeleton organization; Long-term synaptic depression | Synapse; Membrane; Cell junction | Protein binding | *ND* |  |
| **Olfml3** | Olfactomedin-like 3 | Extracellular matrix glycoprotein | *ND* | *ND* | Glial cells | PMID: 30093905 |
| **Zfp420** | Zinc finger protein 420 | Immune system | *ND* | *ND* | Mutation in axonal Charcot-Marie-tooth-disease | PMID: 29341343 |
| **Hspb3** | Heat shock protein 3 | *ND* | Nucleus; Cytoplasm | *ND* |  |  |
| **Penk** | Preproenkephalin | Chemical synaptic transmission; Sensory perception of pain; Glial cell proliferation; Response to morphine | Membrane; Axon; Synapse | Opioid peptide activity | Opioid system in a mice neuropathic pain model | PMID: 30176322 |
| **Rnasek** | Ribonuclease, RNase K | Nucleic acid phosphodiester bond hydrolysis | Membrane | Nuclease activity | *ND* |  |
| **Ddx3y** | Dead (asp-glu-ala-asp) box polypeptide 3, y-linked | Translation machinery | Nucleoplasm; P granule | Nucleic acid binding | *ND* |  |
| **Commd1** | Comm domain containing 1 | Protein transport; Regulation of plasma lipoprotein particle levels | Endosome; membrane; nucleus | Protein binding; Sodium channel inhibitor activity | *ND* |  |
| **Uts2b** | Urotensin 2b | Regulation of blood vessel diameter; Signal transduction | Extracellular region | Hormone activity | *ND* |  |
| **Kdm5d** | Lysine (k)-specific demethylase 5d | Regulation of androgen receptor signaling pathway | Nucleus | DNA binding | *ND* |  |
| **Uty** | Ubiquitously transcribed tetratricopeptide repeat gene, y chromosome | Regulation of gene expression | Membrane | Protein binding | *ND* |  |
| **Eif2s3y** | Eukaryotic translation initiation factor 2, subunit 3, structural gene y-linked | Translation machinery | Eukaryotic translation initiation factor 2 complex | Translation initiation factor activity | *ND* |  |

**Suppl. Table 1**: Functional analysis of differentially expressed INPUT genes. (ND: No data available).

| ***Gene name*** | ***Gene description*** | ***GO Biological Process*** | ***GO Cellular Component*** | ***GO Molecular Function*** | ***Nervous system, nociception related?*** | ***Reference*** |
| --- | --- | --- | --- | --- | --- | --- |
|  | **UP- REGULATED IN FEMALES** | | | | | |
| **Gm527** | Predicted gene 527 | *ND* | *ND* | *ND* | *ND* |  |
| **Pcdha8** | Protocadherin alpha 8 | Cell adhesion; nervous system development | Membrane | Calcium ion binding | May be involved in the establishment and maintenance of specific neuronal connections | PMID: 10380929 |
| **Ccdc17** | Coiled-coil domain containing 17 | *ND* | *ND* | *ND* | *ND* |  |
| **Lime1** | Lck interacting transmembrane adaptor 1 | Regulation of map kinase activity; T and B cell receptor signaling | Membrane | Protein kinase binding | *ND* |  |
| **Map3k1** | Mitogen-activated protein kinase kinase kinase 1 | Protein phosphorylation | *ND* | Protein kinase activity | Inflammatory pain | PMID: 18366630 |
| **Efcab7** | Ef-hand calcium binding domain 7 | Regulates hedgehog | *ND* | Calcium ion binding | *ND* |  |
| **Zmym1** | Zinc finger mym-type containing 1 | Transcription regulation | *ND* | Protein dimerization activity; zinc ion binding | *ND* |  |
| **Mllt10** | Myeloid/lymphoid or mixed-lineage leukemia; translocated to, 10 | Transcription regulation | Nucleus, cytosol | Protein binding | *ND* |  |
| **Gm42878** | Predicted gene 42878 | Protein phosphorylation |  | Protein kinase activity | *ND* |  |
| **Pf4** | Platelet factor 4 | Inflammatory/immune response; adenylate cyclase-activating g protein-coupled receptor signaling pathway | Cytoplasm | Cytokine/chemokine activity | Sickle cell disease | PMID: 2145991 |
| **Necab2** | N-terminal ef-hand calcium binding protein 2 | Positive regulation of ERK1 and ERK2 cascade | Cytoplasm, membrane, axon | Signaling scaffold protein | Inflammatory hypersensitivity; Nerve injury | PMID: 29893745; PMID: 24616509 |
| **Ep400** | E1a binding protein p400 | Chromatin organization | Nucleus | Nucleotide binding | Myelin formation | PMID: 31081019 |
| **Ccdc84** | Coiled-coil domain containing 84 | *ND* | *ND* | *ND* | Cluster headache | PMID: 28074859 |
| **Slc9a2** | Solute carrier family 9 member A2 | Sodium ion transport; protein localization; pH regulation | Membrane | Antiporter activity | *ND* |  |
| **Fbln1** | Fibulin 1 | Negative regulation of erk1 and erk2 cascade | Extracellular matrix | Protein binding | *ND* |  |
| **Klf4** | Kruppel-like factor 4 | Transcription regulation | Nucleus | Nucleic acid binding | Overexpression of KLF4 can regulate neuronal cell cycle proteins and sensitize neurons to NMDA. | PMID: 19041854 |
| **Inpp5d** | Inositol polyphosphate-5-phosphatase d | Signal transduction | Cytoplasm | Protein binding | Alzheimer’s disease | PMID: 30906402; PMID: 28482637 |
| **Dpt** | Dermatopontin | Cell adhesion | Extracellular region | Extracellular matrix structural constituent | Potential role in neuronal functions *in vivo* (zebrafish) | PMID: 23266816 |
| **Hmgxb4** | Hmg box domain containing 4 | Transcription regulation | Nucleus | DNA binding | *ND* |  |
| **Eif2s3x** | Eukaryotic translation initiation factor 2, subunit 3, structural gene x-linked | Formation of translation preinitiation complex | Eukaryotic translation initiation factor 2 complex in cytoplasm | Translation factor activity; RNA binding | *ND* |  |
| **Sfrp4** | Secreted frizzled-related protein 4 | Wnt signaling pathway | Cell | Protein binding | *ND* |  |
| **Rabepk** | Rab9 effector protein with kelch motifs | *ND* | Membrane, endosome | Protein binding | ND |  |
| **Igfbp6** | Insulin-like growth factor binding protein 6 | Regulation of insulin-like growth factor receptor signaling pathway | Cytoplasm | Growth factor binding | Spinal cord injury, peripheral nerve injury | PMID: 27888466; PMID: 9762866 |
| **Zmynd8** | Zinc finger, mynd-type containing 8 | Modulation of excitatory postsynaptic potential | Nucleoplasm, dendrite | Protein binding | Neuronal differentiation | PMID: 20331974 |
| **Mtss1l** | Mtss i-bar domain containing 2 | Membrane organization | Membrane | Actin binding | Membrane curvature and dendritic spine formation, regulation of synapse currents | PMID: 31232686 |
| **Rab11b** | Rab11b, member ras oncogene family | Rab protein signal transduction, protein transport | Synapse, cell, membrane | Non-kinase enzyme | Neuronal endosomal pathways | PMID: 14627637 |
| **Lepr** | Leptin receptor | Regulation of MAPK cascade, signal transduction, protein phosphorylation | Membrane | Transmembrane signaling receptor activity | Neuropathic pain | PMID: 29071294; PMID: 28822786 |
| **Prelp** | Proline arginine-rich end leucine-rich repeat | Cell aging | Extracellular region | Protein binding | Endometriosis | PMID: 28678915 |
| **Gjb6** | Gap junction protein, beta 6 | Gap junction assembly; Cell communication | Cell; Membrane; Gap junction | Gap junction channel activity |  |  |
| **Lbp** | Lipopolysaccharide binding protein | Immune response | Membrane | Signaling receptor binding | Metabolic neuropathy | PMID: 31145897 |
| **Repin1** | Replication initiator 1 | *ND* | Cytosolic ribosome | Nucleic acid binding | *ND* |  |
| **Slc6a13** | Solute carrier family 6 (neurotransmitter transporter, GABA), member 13 | Neurotransmitter transport | Neuron projection; Membrane | Neurotransmitter binding, symporter activity | *ND* |  |
| **Ptgds** | Prostaglandin d2 synthase | Nociception; Sleep regulation; Temperature regulation | Cytoplasm | Prostaglandin-D synthase activity | Nociception | PMID: 9892701 |

| ***Gene name*** | ***Gene description*** | ***GO Biological Process*** | ***GO Cellular Component*** | ***GO Molecular Function*** | ***Nervous system, nociception related?*** | ***Reference*** |
| --- | --- | --- | --- | --- | --- | --- |
|  | **UP- REGULATED IN MALES** | | | | | |
| ***Mapk1ip1*** | Mitogen-activated protein kinase 1 interacting protein 1 | *ND* | Cytoplasm | *ND* | *ND* |  |
| ***Chka*** | Choline kinase alpha | Metabolic process | Cytoplasm | Kinase activity | Neuronal function | PMC2925531 |
| ***Serpinb1b*** | Serine (or cysteine) peptidase inhibitor, clade b, member 1b | *ND* | Cytoplasm | Regulator of enzyme | *ND* |  |
| ***Timp1*** | Tissue inhibitor of metalloproteinase 1 | Signal transduction | *ND* | Regulator of enzyme | Neuropathic pain, nerve injury | PMID: 17033093; PMID: 21782897 |
| ***Ccer2*** | coiled-coil glutamate-rich protein 2 | *ND* | *ND* | *ND* | taxane-induced neuropathic pain | PMC5347688 |
| ***Desi2*** | Desumoylating isopeptidase 2 | Proteolysis | Cytoplasm | Enzyme activity | *ND* |  |
| ***Itga11*** | Integrin alpha 11 | Cell adhesion | Membrane | Collagen binding | *ND* |  |
| ***Apln*** | Apelin | Signal transduction | Extracellular space | Signaling receptor binding | Neuropathic pain, Alzheimer’s disease | PMC5562064; PMID: 29807653 |
| ***Bcat1*** | Branched chain aminotransferase 1, cytosolic | Amino acid byosynthetic and metabolic process | Cytoplasm | Catalytic activity | mTOR pathway in Alzheimer’s disease | PMID: 29802157 |
| ***Chac1*** | Chac, cation transport regulator 1 | Notch signaling pathway, neurogenesis, regulation of protein processing | Cytoplasm | Gamma-glutamylcyclotransferase activity; notch binding | *ND* |  |
| ***Hoxd11*** | Homeobox d11 | Transcription regulation | Nucleus | DNA binding | *ND* |  |
| ***Crym*** | Crystallin, mu | Transcription regulation | Nucleus, cytoplasm | Hormone binding, NADP binding | *ND* |  |
| ***Fgf9*** | Fibroblast growth factor 9 | Wnt signaling; MAPK cascade; cell-cell signaling | Cytoplasm | Fibroblast growth factor receptor binding | Neuron-derived fgf9 | PMID: 19232523; PMID: 31726090 |
| ***Sema6a*** | Sema domain, transmembrane domain (tm), and cytoplasmic domain, (semaphorin) 6a | Neuron migration, axon guidance, regulation of ERK1 and ERK2 cascade | Axon, membrane | *ND* | Axon guidance and neuronal connectivity; burn injury | PMID: 27392094; PMID: 27573516 |
| ***Tbl3*** | Transducin (beta)-like 3 | rRNA processing, post-transcriptional regulation | Pwp2p-containing subcomplex of 90s preribosome | Protein binding, snoRNA binding | *ND* |  |
| ***Esam*** | Endothelial cell-specific adhesion molecule | Cellular protein localization, cell-adhesion | Membrane | *ND* | *ND* |  |
| ***Pde8b*** | Phosphodiesterase 8b | Signal transduction | Hydrolase activity, metal ion binding | Enzyme | Phosphodiesterases in neurodegenerative diseases | PMID: 23129425 |
| ***Ddx4*** | Dead (asp-glu-ala-asp) box polypeptide 4 | Regulation of protein localization, male meiosis | Nucleus, cytoplasm, p granule, ribonucleoprotein complex | RNA binding | *ND* |  |
| ***Nhlh2*** | Nescient helix loop helix 2 | Transcription regulation | Nucleus | Protein binding | Neuronal transcription factor | PMID: 31421827 |
| ***Ccne2*** | Cyclin e2 | Synapsis | Nucleus, cytoplasm | Protein kinase activity |  |  |
| ***Thrb*** | Thyroid hormone receptor beta | Transcription regulation | Nucleus | Protein binding | Neuronal regeneration after brain injury | PMID: 29378220 |
| ***Hoxd10*** | Homeobox d10 | Peripheral nervous system neuron development, transcription regulation | Cytoplasmic ribonucleoprotein granule | | *ND* |  |
| ***Slc16a13*** | Solute carrier family 16 (monocarboxylic acid transporters), member 13 | Transmembrane transport | Membrane | Symporter activity, GABA transporter | *ND* |  |
| ***Serpina3c*** | Serine (or cysteine) peptidase inhibitor, clade a, member 3c | Regulation of peptidase activity, response to cytokine | Extracellular space | Peptidase inhibitor activity | Neuropathic pain | PMID: 25915831 |
| ***Card9*** | Caspase recruitment domain family, member 9 | Positive regulation of interleukin-6 production, regulation of stress-activated MAPK cascade, JNK cascade | Cytoplasm | Protein binding | *ND* |  |
| ***Atf3*** | Activating transcription factor 3 | Transcription regulation | Nucleus | *ND* | Injury marker | PMID: 15066140 |
| ***Gtpbp4*** | GTP binding protein 4 | Ribosome biogenesis, protein stabilization, cell-cell adhesion | Nucleus, cytoplasm, membrane | GTP activity | *ND* |  |
| ***Sez6l*** | Seizure related 6 homolog like | Regulation of protein kinase C signaling, synapse maturation | Membrane, neuronal cell body | Protease substrate in neurons | Burn injury; neuropathic pain | PMID: 27573516; PMID: 25880204 |
| ***Srd5a2*** | Steroid 5 alpha-reductase 2 | Regulation of testosterone; androgen processing |  | Enzyme | *ND* |  |
| ***Kdm5d*** | Lysine (k)-specific demethylase 5d | Regulation of androgen receptor signaling pathway | Nucleus | DNA binding | *ND* |  |
| ***Uty*** | Ubiquitously transcribed tetratricopeptide repeat gene, y chromosome | Regulation of gene expression | Membrane | Protein binding | *ND* |  |
| ***Ddx3y*** | Dead (asp-glu-ala-asp) box polypeptide 3, y-linked | Translation machinery | Nucleoplasm, P granule | Nucleic acid binding | *ND* |  |
| ***Eif2s3y*** | Eukaryotic translation initiation factor 2, subunit 3, structural gene y-linked | Translation machinery | Eukaryotic translation initiation factor 2 complex | Translation initiation factor activity | *ND* |  |

**Suppl. Table 2**: Functional analysis of differentially translated IP mRNAs. (ND: No data available)

**Suppl. File 1 Tab 1A**: Raw transcripts per million for INPUT and respective percentiles.

**Suppl. File 1 Tab 1B**: Quantile normalized transcripts per million for INPUT and respective statistics.

**Suppl. File 1 Tab 2A**: Raw transcripts per million for IP and respective percentiles.

**Suppl. File 1 Tab 2B**: Quantile normalized transcripts per million for IP and respective statistics.

**Suppl. File 2**: Differentially expressed genes in INPUT.

**Suppl. File 3**: Differentially translated mRNAs in IP.

**Suppl. Material and Methods:**

**Animals**

All animal procedures were approved by the Institutional Animal Care and Use Committee of University of Texas at Dallas protocol number 14-04.

Nav1.8Cre/Rosa26^fstrap^ mice**:** Rosa26^fsTRAP^ mice were purchased from The Jackson Laboratory (stock #022367). Transgenic mice expressing Cre recombinase under the control of the Scn10a (Nav1.8) promoter were obtained initially from Professor John Wood (University College London) but are commercially available from Infrafrontier (EMMA ID: 04582). Initial studies demonstrated that the introduction of the Cre recombinase in heterozygous animals does not affect pain behavior, and their DRG neurons have normal electrophysiological properties (1). Nav1.8-cre mice on a C57BL/6J genetic background were maintained and bred at the University of Texas at Dallas. Upon arrival, Rosa26^fsTRAP^ mice were crossed to Nav1.8-cre to generate the Nav1.8-TRAP mice that express a fused EGFP-L10a protein in Nav1.8-positive neurons. TRAP experiments were performed using male only and female only Nav1.8-TRAP littermates 8–12 weeks old. Mice were group housed (4 maximum) in non-environmentally enriched cages with food and water ad libitum on a 12 h light-dark cycle. Room temperature was maintained at 21 ± 2°C. Immunohistochemistry and ELISA assays were performed on C57BL/6J mice. Behavioral experiments were performed on C57BL/6J mice and ICR mice (we did not observe any strain differences).

**TRAP**

Nav1.8-TRAP male and female mice (3 mice per sex were pooled for each of 4 biological replicates) were decapitated under isoflurane anesthesia and DRGs rapidly dissected in ice-cold dissection buffer (1× HBSS (Invitrogen, 14065006), 2.5 mM HEPES-NaOH [pH 7.4], 35 mM Glucose, 5mM MgCl2, 100 μg/ml cycloheximide, 0.2 mg/ml emetine). DRGs were transferred to ice-cold Precellys Tissue homogenizing CKMix tube with lysis buffer (20 mm HEPES, 12 mm MgCl2, 150 mm KCl, 0.5 mm DTT, 100 μg/ml cycloheximide, 20 μg/ml emetine, 80 U/ml SUPERase IN, Promega, 1 μl DNase, and protease inhibitor). The lysate was prepared by homogenizing the samples using Precellys® Minilys Tissue Homogenizer at 10 second intervals for a total of 80 seconds, in the cold room (4°C). Samples were then centrifuged at 2000 × g for 5 min to prepare postnuclear fractions. Next, 1% NP-40 and 30 mM 1,2-dihexanoyl-sn-glycero-3-phosphocholine were added and samples centrifuged at 15,000 × g for 10 min to generate a postmitochondrial fraction. A 200 μl sample of this fraction was saved for use as INPUT (bulk RNA-sequencing), and the remaining was incubated with protein G-coated Dynabeads (Invitrogen) bound to 50 μg anti-GFP antibodies (HtzGFP-19F7 and HtzGFP-19C8, Memorial Sloan Kettering Centre) for 3 h at 4°C with end-over-end mixing. Anti-GFP beads were washed with high salt buffer (20 mm HEPES, 5 mm MgCl2, 350 mm KCl, 1% NP-40, 0.5 mm DTT, and 100 μg/ml cycloheximide), and RNA was eluted from all samples using the Direct-zol kit (Zymo Research) according to the manufacturer’s instructions. RNA yield was quantified using a Nanodrop system (Thermo Fisher Scientific), and RNA quality was determined by fragment Analyzer (Advanced Analytical Technologies).

**Immunohistochemistry**

Animals were anesthetized with isoflurane (4%) and euthanized by decapitation and tissues were flash frozen in OCT on dry ice. Sections of DRG (20 μm) and brain (20 μm) were mounted onto SuperFrost Plus slides (Thermo Fisher Scientific) and immediately fixed in ice-cold formalin (10%) for 15 min followed by dehydration in 50% ethanol, 70% ethanol and 100% ethanol at room temperature for 5 min each. Tissues were briefly air dried and boundaries were drawn around each section using a hydrophobic pen (ImmEdge H-4000). Once dry, the sections were blocked for at least 1 h in 10% normal goat serum with 0.3% TX-100. DRG slices were stained with peripherin, NeuN and Ptgds overnight at 4°C and brain slices were stained with NeuN and Ptgds for 2 h at room temperature. The peripherin antibody (P5117) was obtained from Sigma-Aldrich, NeuN (MAB377) was obtained from Millipore Sigma and Ptgds (ab182141) from Abcam. Sections were then washed and incubated with respective Alexa Fluor secondary antibodies for 1 h at room temperature. Sections were washed, air dried, and then coverslipped with Prolong Gold Antifade reagent (Fisher Scientific; P36930).

DRG images were taken using an Olympus FluoView 1200 confocal microscope, using the same settings for all images. Analysis of DRG images was done using ImageJ version 1.48 (National Institutes of Health, Bethesda, MD). The values plotted on Figure 5E were obtained by calculating the corrected total cell fluorescence (CTCF) using the following formula: CTCF= Integrated Density – (Area of selected cell X Mean fluorescence of background readings).

Brain images were taken using Olympus vs120 virtual slide microscope, using the same settings for all images. Analysis of brain images was performed using Olympus cellSens software. An ROI of the same size was placed over 100 randomly selected cortical neurons per section to measure the mean gray intensity value. 3 sections were analyzed per animal. Background fluorescence was measured similarly using a negative control that was only exposed to blocking solution and secondary antibodies (no primary). After subtracting background values of negative control, we averaged all intensity values (first per slice and second per animal) to obtain the values plotted on Suppl. Figure 1.

**Ptgds validation and Estrous cycle assessment**

A separate TRAP experiment was conducted where we validated the expression of Ptgds bound to ribosome. We followed the same protocol as described above (TRAP) with two exceptions: we used only one mouse per replicate, and we tracked the female estrous cycle. The stage of the cycle was determined by vaginal lavage followed by cytological evaluation according to a previous published protocol (2).

**ELISA assay**

PGD_2_ levels in the DRG were evaluated using Prostaglandin D2-MOX Express ELISA Kit (Cayman, 500151) following the manufacturer’s instructions. The same number of DRGs (collected from all levels cervical, thoracic and lumbar) were used per sample in the ELISA assay.

**Injections**

PGE_2_ (Cayman, 14010) was diluted in sterile PBS and injected with a volume of 25 μl via a 30.5-gauge needle and given intraplantarly.

AT-56 (Tocris, 3531) was diluted in 10% DMSO and 30% cyclodextrine and administered intraperitoneally.

**Behavior testing**

All behavioral experiments were performed between 8:00 A.M. and 6:00 P.M. Facial grimacing was evaluated using the Mouse Grimace Scale (MGS) as described previously (3). Mechanical paw withdrawal thresholds were measured using the up-down method (4) with calibrated Von Frey filaments (Stoelting).

**Library generation and sequencing**

After purifying the RNA, cDNA libraries were prepared with total RNA Gold library preparation (with ribosomal RNA depletion) for all samples according to the manufacturer’s instructions (Illumina). Quality control was performed for RNA extraction and cDNA library preparation steps with Qubit (Invitrogen) and High Sensitivity NGS fragment analysis kit on the Fragment Analyzer (Agilent Technologies). After standardizing the amount of cDNA per sample, the libraries were sequenced on Illumina NextSeq500 sequencing machine with 75-bp single-end reads. mRNA library preparation and sequencing was done at the Genome Center in the University of Texas at Dallas Research Core Facilities.

**Mapping and TPM quantification:**

RNA-seq read files (fastq files) were checked for quality by FastQC (Babraham Bioinformatics, <https://www.bioinformatics.babraham.ac.uk/projects/fastqc/>) and read trimming was done based on the Phred score and per-base sequence content. Trimmed Reads were then mapped against the reference genome and transcriptome (Gencode vM16 and GRCm38.p5) using STAR v2.2.1 (5). Relative abundances in Transcripts Per Million (TPM) for every gene of every sample was quantified by stringtie v1.3.5 (6). Non-coding genes and mitochondrial genes were removed from the analysis (based on Gencode annotation) and the TPMs for coding genes were re-normalized to sum to 1 million.

**Order statistics and re-normalization of expression data**

In order to identify a set of consistently expressed genes in the transcriptome (INPUT) samples, percentile ranks were calculated on TPMs for each coding gene for each sample. We conservatively chose 15,072 genes that were above the 30th percentile in each INPUT sample, for at least one sex, to be in the set of genes considered consistently detected in the transcriptome. Quantile normalization was then performed based on the set of all coding genes.

The IP (translatome) analysis was only performed for the 15,072 consistently transcriptome-expressed genes. In order to identify a set of consistently expressed genes in the translatome (IP) samples, percentile ranks were calculated on TPMs for each of the 15,072 consistently transcriptome-expressed coding genes for each sample. We then chose 12,542 genes out of those 15,072 genes to be consistently detected in the translatome based on whether their expression was on or above the 15th percentile (out of 15,072 genes) in each IP sample, for at least one sex.

The percentile thresholds for choosing consistently transcriptome-expressed and translatome-expressed genes were conservatively estimated by identifying thresholds that would eliminate genes with consistently low or no detected reads.

**Differential expression analysis**

We first calculated the log2-fold change (based on median TPMs) for each consistently transcriptome-expressed coding gene in the INPUT, and for each consistently translatome-expressed coding gene in the IP. It is calculated as follows :

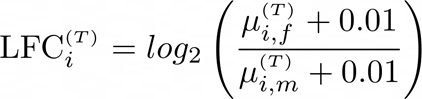

where for gene i, LFC(T)i is the log2-fold change, μ(T)i,f and μ(T)i,m are the median TPMs in females and males respectively, with 0.01 as the smoothing factor.

We used strictly standardized mean difference (SSMD) (6, 7) to discover genes with systematically altered expression percentile ranks between males and females. SSMD is the difference of means controlled by the variance of the sample measurements. We used SSMD as a measure of effect size since it is appropriate for smaller sample sizes while simultaneously controlling for within-group variability. It is calculated as follows :

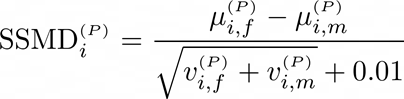

where for gene i, SSMD(P)i is the strictly standardized mean difference, with μ(P)i,f , v(P)i,f and μ(P)i,m , v(P)i,f are the means and variances of gene TPM percentile ranks in females and males respectively, under the assumption that covariance is 0, and with 0.01 as the smoothing factor.

For calculating the overlap in distribution between the qTPMs, in animals sampled from different sexes, we further calculated Bhattacharyya distance (8), which is used to calculate the amount of overlap in the area under the curve of the two sample distributions (corresponding to each sex) in order to identify the best candidates for the DE gene set. Unlike SSMD, BD does not make assumptions of equal variance in the two compared samples, and thus, is useful for comparing distributions of gene relative abundance (in TPMs). It is calculated as follows:

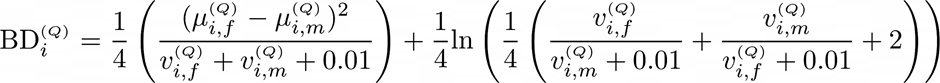

where for gene i, BD(Q)i is the Bhattacharyya Distance over gene qTPMs, with μ(Q)i,f , v(Q)i,f and μ(Q)i,m , v(Q)i,f are the means and variances of gene qTPMs in females and males respectively, and with 0.01 as the smoothing factor. The Bhattacharyya coefficient BC(Q)i ranges between 0 (for totally non-overlapping distributions) and 1 (for completely identical distributions) and is derived from the Bhattacharyya distance as follows:

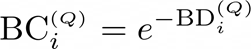

In our analysis, we used a modified form of the Bhattacharyya coefficient that ranges between 0 (for completely identical distributions) and +1 or -1 (for totally non-overlapping distributions, sign defined by the log-fold change value). It is calculated as follows:

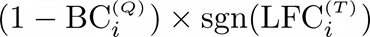

#### Motif analysis

For each gene in the up-regulated translation gene list (IP fraction), we created one 5′ UTR and 3’ UTR sequence per gene. For each gene, this was done by first extracting the respective UTR sequences from all isoforms from BioMart (7). Isoforms with UTRs <20 bases, or with sequence that is part of an ORF in another isoform were discarded. The remaining isoforms were collapsed into a single sequence in order of their location in the genome, with overlapping regions present exactly once in the sequence. The motif finding tool MEME (8) was used to identify motifs.

**Network of interactions**

We used STRING database (9) and Cytoscape (10) to generate Figure 3G and visualize interactions between genes differentially expressed and DRG enriched genes.

**Single cell data**

Single cell mouse DRG sequencing data from previously published work (11) was used to generate Figure 5B. Seurat package 2.2.1 (12) was used to cluster the single-cell data and visualization (t-SNE) (13).

#### Statistics

Statistical analysis for behavior, image quantification and ELISA assay was done in GraphPad Prism 8. Data visualization was done in Python (version 3.7 with Anaconda distribution). All data are represented as mean ± SEM. Single comparisons were performed using Student’s t test, and multiple comparisons were performed using a one-way or two-way ANOVA with post hoc tests for between-group comparisons. Statistical results can be found in the figure legends.

Coding for bioinformatics analysis and data visualization was done in Python (version 3.7 with Anaconda distribution).  Sequencing data is deposited in GEO accession number.

Figure 1 and other schematic drawings were generated using Biorender (BioRender.com).

1.	Stirling LC, Forlani G, Baker MD, Wood JN, Matthews EA, Dickenson AH, et al. (2005): Nociceptor-specific gene deletion using heterozygous NaV1. 8-Cre recombinase mice. *Pain*. 113:27-36.

2.	McLean AC, Valenzuela N, Fai S, Bennett SAL (2012): Performing vaginal lavage, crystal violet staining, and vaginal cytological evaluation for mouse estrous cycle staging identification. *J Vis Exp*.e4389-e4389.

3.	Langford DJ, Bailey AL, Chanda ML, Clarke SE, Drummond TE, Echols S, et al. (2010): Coding of facial expressions of pain in the laboratory mouse. *Nat Methods*. 7:447-449.

4.	Chaplan SR, Bach FW, Pogrel JW, Chung JM, Yaksh TL (1994): Quantitative assessment of tactile allodynia in the rat paw. *J Neurosci Methods*. 53:55-63.

5.	Dobin A, Davis CA, Schlesinger F, Drenkow J, Zaleski C, Jha S, et al. (2013): STAR: ultrafast universal RNA-seq aligner. *Bioinformatics*. 29:15-21.

6.	Pertea M, Pertea GM, Antonescu CM, Chang T-C, Mendell JT, Salzberg SL (2015): StringTie enables improved reconstruction of a transcriptome from RNA-seq reads. *Nature biotechnology*. 33:290-295.

7.	Kinsella RJ, Kähäri A, Haider S, Zamora J, Proctor G, Spudich G, et al. (2011): Ensembl BioMarts: a hub for data retrieval across taxonomic space. *Database (Oxford)*. 2011:bar030.

8.	Bailey TL, Boden M, Buske FA, Frith M, Grant CE, Clementi L, et al. (2009): MEME SUITE: tools for motif discovery and searching. *Nucleic Acids Res*. 37:W202-208.

9.	Szklarczyk D, Gable AL, Lyon D, Junge A, Wyder S, Huerta-Cepas J, et al. (2019): STRING v11: protein–protein association networks with increased coverage, supporting functional discovery in genome-wide experimental datasets. *Nucleic acids research*. 47:D607-D613.

10.	Shannon P, Markiel A, Ozier O, Baliga NS, Wang JT, Ramage D, et al. (2003): Cytoscape: a software environment for integrated models of biomolecular interaction networks. *Genome research*. 13:2498-2504.

11.	Li CL, Li KC, Wu D, Chen Y, Luo H, Zhao JR, et al. (2016): Somatosensory neuron types identified by high-coverage single-cell RNA-sequencing and functional heterogeneity. *Cell Res*. 26:967.

12.	Butler A, Hoffman P, Smibert P, Papalexi E, Satija R (2018): Integrating single-cell transcriptomic data across different conditions, technologies, and species. *Nat Biotechnol*. 36:411-420.

13.	Maaten Lvd, Hinton G (2008): Visualizing data using t-SNE. *Journal of machine learning research*. 9:2579-2605.
